# LSD flattens the hierarchy of directed information flow in fast whole-brain dynamics

**DOI:** 10.1101/2024.04.25.591130

**Authors:** Kenneth Shinozuka, Prejaas K.B. Tewarie, Andrea Luppi, Christopher Lynn, Leor Roseman, Suresh Muthukumaraswamy, David J. Nutt, Robin Carhart-Harris, Gustavo Deco, Morten L. Kringelbach

## Abstract

Psychedelics are serotonergic drugs that profoundly alter consciousness, yet their neural mechanisms are not fully understood. A popular theory, RElaxed Beliefs Under pSychedelics (REBUS), posits that psychedelics flatten the hierarchy of information flow in the brain. Here, we investigate hierarchy based on the imbalance between sending and receiving brain signals, as determined by directed functional connectivity. We measure directed functional hierarchy in a magnetoencephalography (MEG) dataset of 16 healthy human participants who were administered a psychedelic dose (75 micrograms, intravenous) of lysergic acid diethylamide (LSD) under four different conditions. LSD diminishes the asymmetry of directed connectivity when averaged across time. Additionally, we demonstrate that machine learning classifiers distinguish between LSD and placebo more accurately when trained on one of our hierarchy metrics than when trained on traditional measures of functional connectivity. Taken together, these results indicate that LSD weakens the hierarchy of directed connectivity in the brain by increasing the balance between senders and receivers of neural signals.

## Section 1. Introduction

Classic psychedelic compounds, such as psilocybin and lysergic acid diethylamide (LSD), are serotonergic drugs that have recently been investigated for their ability to treat a wide range of psychiatric disorders, including treatment-resistant depression, end-of-life anxiety, PTSD, and addiction (Cavarra et al., 2022). They are specifically known to alter conscious experience through their agonist action at the 5-HT_2A_ receptor (Gonzalez-Maeso & Sealfon, 2009; López-Giménez & González-Maeso, 2018; Nichols, 2016; Vollenweider et al., 1998). In the last decade, neuroscientists have used functional magnetic resonance imaging (fMRI), magnetoencephalography (MEG), electroencephalography (EEG), and other modalities to explore the effects of psychedelics on the brain, both in healthy participants (e.g. Carhart-Harris et al., 2016) and clinical populations (e.g. Carhart-Harris et al., (2017); Doss et al., (2021)).

A recent theory, RElaxed Beliefs Under pSychedelics (REBUS), claims that psychedelics “flatten” hierarchical information processing in the brain (R. L. Carhart-Harris & Friston, 2019). Based on the predictive coding model of the brain, REBUS claims that strongly-weighted expectations, or priors, encoded in higher-order, associative brain regions constrain the bottom-up propagation of sensory information from lower-order areas (Holmes & Nolte, 2019; Knill & Pouget, 2004; Seth & Friston, 2016). REBUS hypothesises that psychedelics decrease the weighting or strength of activity instantiating these priors, but it is challenging to use techniques such as fMRI and MEG to probe the encoding of these priors in the brain and how this encoding may be perturbed under psychedelics.

With fMRI and EEG/MEG, it is more feasible to test one implication of REBUS’ hypothesis, namely that psychedelics diminish top-down information flow while enhancing bottom-up information flow. For instance, based on fMRI data, Girn and colleagues identified a principal gradient of functional connectivity (FC) spanning higher-order and lower-order brain regions and observed that LSD and psilocybin cause this gradient to contract (Girn et al., 2022), a finding that has since been replicated with the psychedelic N,N-dimethyltryptamine (DMT) (Timmermann et al., 2023). Other research has extracted “backward” (top-down) and “forward” (bottom-up) travelling waves from EEG data and demonstrated that DMT suppresses the former while elevating the latter (Alamia et al., 2020; Timmermann et al., 2023). Finally, psilocybin decreases directed information flow, as measured with Granger causality in MEG data, towards frontal regions of the brain and away from more posterior areas (Barnett et al., 2020).

While the principal gradient analysis captures undirected connectivity, both travelling waves and Granger causality are directed measures. Indeed, it is intuitive to investigate hierarchies based on directional relationships; for example, nodes at the top of a hierarchy may exert (top-down) directed control over the activity in nodes at lower levels. The same interaction cannot be inverted – subordinate areas do not control higher-order areas – so the information flow is asymmetric. We could therefore define hierarchy as an imbalance or asymmetry in the directionality of information flow.

In particular, we can distinguish three different properties of hierarchies: irreversibility, hierarchical coherence, and hierarchical inhomogeneity. To measure irreversibility, we will apply a technique known as INSIDEOUT, which computes the difference between “forwards” directed connectivity (e.g., from a region *A* to a region *B*) and time-reversed directed connectivity (e.g., from *B* to *A*), averaged across all *pairs* of regions (Deco et al., 2022). Hierarchical coherence and inhomogeneity are the mean and standard deviation of this asymmetry when it is measured at *each* region. We quantified these three characteristics of hierarchy in previously-acquired MEG data of 16 healthy human participants who were administered a psychoactive dose of LSD (Carhart-Harris et al., 2016).

## Section 2. Methods

### Section 2.1. Data acquisition

The data was acquired in a previous study (Carhart-Harris et al., 2016), and the experimental protocol is described in detail in the supplementary materials therein. Each participant underwent two scanning sessions, one in which they received placebo and another in which they received 75 μg of intravenously administered LSD. After a 60-minute period of acclimatisation, participants were scanned with magnetic resonance imaging (MRI) for about 60 minutes; the results of the MRI scans are not discussed in this analysis but have been discussed extensively in other publications (Atasoy et al., 2017; Girn et al., 2022; Jobst et al., 2021; Kaelen et al., 2016, 2017; Lebedev et al., 2016; Luppi et al., 2021; Roseman et al., 2016; Singleton et al., 2022; Tagliazucchi et al., 2016; Varley et al., 2020). Then, approximately 165 minutes after the LSD was administered, the participants were recorded with a CTF 275-gradiometer MEG, though four of the sensors were turned off because of excessive sensor noise. During the MEG recording, participants were subjected to four conditions: eyes-open resting-state (referred to in the paper as “Open”), eyes-closed resting-state (“Closed”), listening to music with eyes closed (“Music”), and watching a silent nature documentary (“Video”). Each scan lasted approximately seven minutes. There were two scans associated with each of the Open and Closed conditions and one scan for each of the Music and Video conditions. We randomly selected one Open and one Closed scan per subject.

### Section 2.2. Data preprocessing and source reconstruction

Preprocessing of the data has also been described elsewhere (Mediano et al., 2024). Data was recorded at 600 Hz, high- and low-pass filtered at 1 Hz and 100 Hz, respectively, segmented into epochs of 2 seconds in length, and, in this study, downsampled to 200 Hz. Epochs with large ocular, muscle, cardiac, and line noise artefacts were removed first manually and then automatically with various algorithms, including ICA. Of the original 20 participants, three were excluded from the data analysis because they either failed to complete both scanning sessions or because their movement artefacts were too large. Additionally, one participant failed to complete all four conditions; they were missing data from the music condition. Therefore, we analysed data from 16 participants.

A linearly constrained minimum variance (LCMV) beamformer was applied to project the preprocessed sensor-space data onto neural sources, which are 90 regions of interest (ROIs) from the AAL parcellation (Van Veen et al., 1997). In the main text, we present the results of our broadband analysis, and we report the frequency-specific analysis in the supplementary materials. For the latter, we bandpass filtered the data into four frequency bands of interest: theta (4-8 Hz), alpha (8-13 Hz), beta (13-30 Hz), and gamma (30-48 Hz) (note that we filtered the data after beamforming). We omitted the delta band because the data was segmented into 2-second epochs, so one epoch could be the same length as a single, low-frequency delta cycle. We applied symmetric orthogonalisation to remove spurious zero-lag correlations from the filtered data (Colclough et al., 2015), and then we obtained the amplitude envelopes of the signal from the Hilbert transform.

### Section 2.3. Irreversibility of hierarchy: INSIDEOUT

Here, we used a technique called INSIDEOUT to measure irreversibility: the squared difference between the cross-correlations of the forwards and reversed amplitude envelope timeseries of all pairs of brain regions (Tewarie et al., 2023).

In general, the cross-correlation between two timeseries *x* and *y* is the correlation between *x* and *y* when *y* is shifted in time relative to *x* by a constant interval or “lag” *τ*. Therefore, for a particular *τ*, the cross-correlations of the forwards and time-reversed timeseries are:

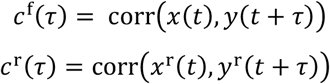

If the system is reversible, then the delay *τ* will not induce temporal asymmetry in the correlation. Because it takes time for signals to travel from one region of the brain to another, and therefore for those regions to become correlated with each other, the value of *τ* that maximises the cross-correlation in MEG may capture the speed of signal propagation for postsynaptic potentials (Mizuno-Matsumoto et al., 2002, 2005). Therefore, for the correct *τ*, the difference in forwards and reversed cross-correlations may indicate the irreversibility of the transmission of postsynaptic potentials in the brain.

We computed the cross-correlations between the amplitude envelopes of all pairs of the 90 brain regions, for all possible values of *τ*. To calculate the cross-correlations, we used the function `xcorr` in MATLAB 2020b. (Note that, before running `xcorr`, we first demeaned the amplitude envelopes.) We measured cross-correlations separately for each epoch, and there were 400 samples in each epoch; thus, there were 399 possible values of *τ*, if *τ* is restricted to positive values only.

We measured *c*^*f*^(*τ*) and *c*^*r*^(*τ*) on the amplitude envelopes of all pairs of the 90 brain regions (both cortical and subcortical, but not cerebellar) in the AAL parcellation, resulting in 90 x 90 *C*^*f*^(*τ*) and *C*^*r*^(*τ*) matrices for each epoch. We then calculated the irreversibility matrix, (*τ*), as the squared difference between each element of *C*^*f*^(*τ*) and *C*^*r*^(*τ*):

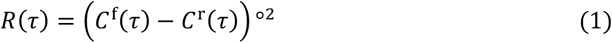

where °^2^ denotes element-wise squaring of *C*^*f*^(*τ*) − *C*^*r*^(*τ*).

We also calculated the irreversibility *R*_*region*_(*τ*) for each region:

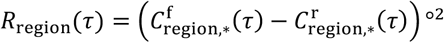

where 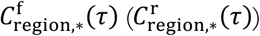 denotes the row of *C*^*f*^(*τ*) (*C*^*r*^(*τ*)) that contains the forward (time-reversed) cross-correlations between the given region and all other regions.

In order to reduce noise, we thresholded (*τ*) and *R*_*region*_(*τ*) by their 95^th^ percentile values, resulting in 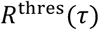 and 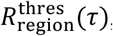, respectively. The choice of the 95^th^ percentile was arbitrary. We also reran our computation of irreversibility without thresholding. The results agreed with the thresholded analysis; we found significant differences in irreversibility between LSD and placebo across all four of the conditions (results not shown).

Finally, we took the mean of *R*^*thres*^(*τ*) and 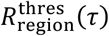, resulting in scalar values of irreversibility *r*(*τ*) and *r*_*region*_(*τ*) for each epoch, respectively. We then averaged these values across all epochs, yielding the final value of irreversibility.

Irreversibility is equivalent to directional asymmetry; that is, the irreversibility of the connection between a timeseries *x* and *y* is the same as the squared difference between the connectivity from *x* to *y* and the connectivity from *y* to *x*. In other words,

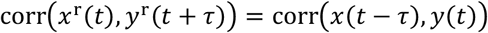

That is, the cross-correlation of the reversed timeseries, with one timeseries shifted forward by *τ*, is equivalent to the cross-correlation of the original timeseries, with one timeseries shifted backwards by *τ*. Furthermore, *corr*(*x*(*t* − *τ*), *y*(*t*)) is simply the correlation of *x* and *y* when *y* is shifted forward in time relative to *x*, i.e. the cross-correlation of *y* with respect to *x*. Therefore, irreversibility is equal to the squared difference between the cross-correlation of *x* with respect to *y* and the cross-correlation of *y* with respect to *x*. For this reason, we equate irreversibility with directional asymmetry.

For this reason, *C*^*r*^(*τ*) is simply equal to the transpose of *C*^*f*^(*τ*). Therefore, **Equation 1** could be re-expressed as:

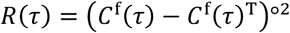

### Section 2.3.1. Choosing τ and statistical testing

The value of a cross-correlation, and therefore the value of irreversibility, depends on the choice of *τ*. We applied three different methods for choosing *τ*. In the first one, we did not identify a specific, optimal *τ*, but instead we computed irreversibility across all possible values of *τ*. We then integrated the area under the irreversibility vs. *τ* curve to obtain a measure of “total” irreversibility. In the second approach, we averaged the irreversibility results across both drugs, all conditions, and all participants (in other words, across all datasets). Then, we selected the *τ* that yielded the highest mean irreversibility. Finally, in the third approach, we used stratified cross-validation and permutation testing to identify the *τ* that was associated with the greatest difference in irreversibility between placebo and LSD.

In the main text, we report the results of Approach 3 since it was the method that aligned most with our aims; it identified the strongest differences in irreversibility between the placebo and LSD datasets. We discuss the results of the other approaches in the Supplementary Materials. For all subsequent measures – hierarchical coherence and inhomogeneity, dynamical analysis, etc. – we used the *τ* derived from Approach 3.

In Approach 1, we applied cluster-based permutation testing separately to each condition in order to determine clusters of *τ* for which irreversibility is significantly different between LSD and placebo (Maris & Oostenveld, 2007). In Approaches 2 and 3, once *τ* was selected, we performed a two-way repeated-measures Analysis of Variance (ANOVA) in which the two within-subject factors were drug and condition, and then we conducted post-hoc Tukey tests if needed. We used a battery of tests to determine whether the distribution of irreversibility at the selected *τ* met the condition of approximate normality (Öner & Kocakoç, 2017). For the *τ* identified in both Approaches 2 and 3, the distribution of irreversibility was approximately normal according to at least two tests. While the placebo results passed almost all normality tests, the LSD results failed the majority (but not all) of the normality tests, but only in the Open condition.

We conducted ANOVAs for all other measures of interest, such as hierarchical coherence and inhomogeneity. Whenever we compared LSD and placebo in specific regions, e.g., for regional irreversibility, we used a three-way repeated measures ANOVA, in which region was a third within-subject factor.

### Section 2.4. Hierarchical coherence and inhomogeneity

Time-delayed correlations define an incoming (e.g., *B* → *A*) and an outgoing (e.g., *A* → *B*) component in each connection in the brain. Irreversibility identifies the *connections* with the highest squared difference between their incoming and outgoing components. Here, we measured a separate aspect of hierarchy: the difference between the magnitudes of the strongest incoming and outgoing connections *at each region*. If the strongest incoming connections are greater in magnitude than the strongest outgoing connections, or vice versa, then the region exhibits imbalanced connectivity.

In *Section 2.3*, we defined *C*^*f*^(*τ*) and *C*^*r*^(*τ*) in each epoch of data. Here, we concatenated 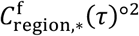 and 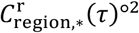 into a single vector; note that these quantities represent the magnitude of the incoming and outgoing connections at each region:

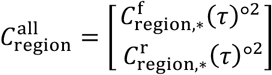

We then thresholded this vector by its 95^th^ percentile, made all outgoing connections that survived thresholding negative, and took the mean of the resulting vector. We refer to this quantity as *A*_*region*_, the incoming-outgoing asymmetry at each region. Then, we average *A*_*region*_ across epochs. The mean of this quantity across regions is the hierarchical coherence, and the standard deviation is the hierarchical inhomogeneity.

### Section 2.5. Dynamical analysis

We wanted to understand how the functional hierarchy evolves over time, both under placebo and under LSD. To do so, we performed a recurrence network analysis on the irreversibility, hierarchical coherence, and hierarchical inhomogeneity. This analysis identifies dynamical patterns that each metric recurrently exhibits. The key idea is to reconstruct the state space of the dynamics by performing a time-delay embedding of the corresponding timeseries, which shifts the timeseries by a series of time-delays.

First, we conducted a sliding-window analysis in which we computed the irreversibility, hierarchical coherence, and hierarchical inhomogeneity in 1-second windows of data, with 80% overlap between windows. This results in three 1D timeseries, one for each hierarchy metric. Secondly, we performed a time-delay embedding of each timeseries; that is, we embedded each timeseries in a lower-dimensional phase space, in which each dimension *d* represents the centrality at lag (*d* – 1) * *τ*_RNoptimal_. This phase space captures temporal dependencies in the centrality timeseries, which is assumed to have regular patterns that re-occur at multiples of *τ*_RNoptimal_. (Note that *τ*_RNoptimal_ is distinct from the *τ* described elsewhere in this paper.) We determined *τ*_RNoptimal_ by computing the mutual information between the timeseries at time *t* and the timeseries at time *t* + 1 for values of *τ*_RN_ ranging from 1 sample to half the length of the timeseries (Wallot & Mønster, 2018). We then selected *τ*_RNoptimal_ as the value of *τ*_RN_ that corresponded to the first minimum of mutual information; at this point, the timeseries shifted by *τ*_RN_ is similar enough to the unshifted timeseries that the embedding reflects genuine temporal dependencies in the data, but not so similar that the shifted timeseries is effectively redundant with the original timeseries.

Next, we determined the dimensionality *m*_optimal_ of the phase space based on the False Nearest Neighbours (FNN) criterion (Wallot & Mønster, 2018). When *m* is too low, many points in the phase space will cluster next to each other, resulting in “false neighbours” that yield false estimates of similarity. For values of *m* ranging from 1 to 10, we calculated the distance between neighbouring points in dimension *m* and again in dimension *m* + 1. If the distance in dimension *m* + 1 was more than twice as high as the distance in dimension *m*, then the points were considered false neighbours in dimension *m*. We selected *m*_optimal_ as the value of *m* that minimised the number of false neighbours. Analyses for determining *τ*_RNoptimal_ and *m*_optimal_ were conducted using the Cross Recurrence Plot (CRP) toolbox provided by the Toolboxes for Complex Systems (TOCSY) (Marwan et al., 2007).

Having optimised *τ* and *m*, we were prepared to construct the recurrence network, which identifies centrality dynamics that the brain recurrently exhibits. The recurrence network *R* is a *T* x *T* matrix with the following elements (Varley & Sporns, 2021):

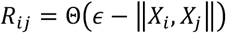

where *X*_*i*_, *X*_*j*_ are timepoints *i* and *j* in the timeseries *X*, ε is a threshold on their distance in phase space, and Θ is the Heaviside function, which transforms ∈ − ‖*X*_*i*_, *X*_*j*_‖ into 1 if it is greater than 0, and 0, if it is less than zero. In other words, if the distance between two timepoints in the phase space is greater than the threshold, then those timepoints are not connected in the recurrence network, but if the distance is less than the threshold, then they are connected. To obtain the distance threshold ε, we used Eroglu’s method (Eroglu et al., 2014). That is, incrementally increasing binary thresholds are applied to the matrix until the matrix becomes connected. Note that a separate recurrence network is obtained for each of the three hierarchy metrics.

Finally, once the recurrence network was constructed, we assessed the recurrency of the hierarchy metrics by measuring the recurrence rate, or edge density, of the network (see *Section 2.7*). If the edge density is high, then the distance between points in the phase space is low, indicating that the dynamics of the corresponding hierarchy metric are similar over time.

### Section 2.6. Random forest classifier

We wished to determine whether irreversibility, hierarchical coherence, and/or hierarchical inhomogeneity distinguish LSD and placebo more than other metrics of FC: undirected FC and (forward) cross-correlations. To obtain undirected FC, we measured the Pearson correlation coefficient between the amplitude envelopes of the timeseries of all 90 regions. We computed (forward) cross-correlations by taking the cross-correlation across all pairs of regions at the value of *τ* identified in Approach 3 in *Section 2.3.1* (*τ*_optimal_ = 0.225 seconds). We then took the mean of each quantity across regions.

We then trained a random forest algorithm with bootstrap aggregation (or “bagging”) to classify LSD and placebo based on undirected FC, cross-correlations, and the three hierarchy metrics. In this method, many random subsets of training data are selected with replacement (Breiman, 1996). These subsets are then used to train a classification tree, which optimises a series of decision points, or thresholds, for splitting the values of the training features in order to estimate the correct label of the training data. In our case, there were two labels, “placebo” or “drug,” and four features, which corresponded to the four conditions. For each classification tree, random forest classifiers choose a random subset of features at each decision point and then identify the feature that optimally splits the training data (Breiman, 2001). Once trained, each classification tree produces a “vote” on the correct label of the test data. By default, `fitcensemble` trains 100 classification trees and then outputs the majority vote of all the trees. We executed `fitcensemble` 100 times, each time with a different training and test set. In each iteration, 80% of the data was partitioned into the training set, and the remaining 20% was partitioned into the test set.

To evaluate the performance of the classifiers, we computed the area under the Receiver Operating Characteristic (ROC) curve. We then performed Wilcoxon signed-rank tests to assess significant differences between the area under the ROC curves for the classifier trained on regional irreversibility and for the classifiers trained on the other FC metrics.

### Section 2.7. Correlation between serotonin receptor expression and regional irreversibility

As mentioned above (*Section 2.3*), we calculated irreversibility within each region of the brain, in addition to computing it across the entire brain. Since the binding of psychedelics to the 5-HT_2A_ (serotonin-2A) receptor is known to meditate their hallucinogenic effects, we measured the correlation between regional irreversibility and 5-HT_2A_ receptor expression. To quantify spatial patterns of 5-HT_2A_ receptor expression, we used three existing PET maps (Beliveau et al., 2017; Savli et al., 2012; Talbot et al., 2012) and one average of those maps (Hansen et al., 2022). We hypothesised that 5-HT_2A_ receptor expression would be more strongly associated with regional irreversibility than that of other serotonin receptors, which may not be essential for the characteristic behavioral effects of psychedelics. Therefore, we also correlated regional irreversibility with 5-HT_1A_ and 5-HT_1B_ receptor expression, using existing PET maps (Beliveau et al., 2017; Gallezot et al., 2010; Hansen et al., 2022; Savli et al., 2012).

To determine the serotonin receptor expression in each AAL region, we averaged serotonin expression across all of the voxels within each region, weighted by the probability that each voxel is in grey matter (Blaiotta et al., 2016). We measured the significance of the correlations by shuffling the serotonin receptor maps 1,000 times in a way that preserved their spatial autocorrelation (Burt et al., 2020). In this method, shuffled maps are smoothed using a kernel-weighted sum of the *𝒦* nearest neighbours of each region, in which the kernel decays exponentially with distance. The algorithm identifies the value of *𝒦* that minimizes the difference between the variogram (variance as a function of distance, which quantifies spatial autocorrelation) of the shuffled and empirical maps. The empirical map is then compared to a null distribution formed from the shuffled maps that were smoothed with the optimal value of *𝒦*.

## Section 3. Results

### Section 3.1. LSD reduces hierarchical irreversibility across multiple conditions

Here, we defined irreversibility as the squared difference between time-delayed correlations of forwards and reversed timeseries. We first calculated irreversibility across all possible values of the time-delay, or *τ*. At all *τ* except for the two or three smallest values of *τ*, placebo was significantly more irreversible in each of the four conditions: Open, Closed, Music, Video (*p*_Open_ = 0.0002, *p*_Closed_ = 0.0001, *p*_Music_ = 0.0001, *p*_Video_ = 0.001) (**Supplementary Figure 1**). We integrated the *τ* vs irreversibility function to obtain a measure of “total irreversibility” across all *τ*, which LSD significantly decreased across conditions (*p*_Open_ < 0.0001, *p*_Closed_ < 0.0001, *p*_Music_ < 0.0001, *p*_Video_ = 0.0023) (**Supplementary Figure 2**). We also found that the *τ* maximising irreversibility was significantly lower on LSD in all four conditions (*p*_Open_ = 0.0009, *p*_Closed_ = 0.0044, *p*_Music_ = 0.0214, *p*_Video_ = 0.0012) (**Supplementary Figure 3**). Intriguingly, across all four conditions, the variance across participants of the *τ* that maximised irreversibility was much greater in LSD than in placebo; for instance, this value of *τ* ranged from 0.025 to 0.08 seconds on LSD in the Open condition, whereas it was 0.08 seconds for every participant on placebo in the same condition. We used two other methods to identify a single value of *τ* for computing irreversibility. In the first, we determined the *τ* that maximised irreversibility across all conditions, drugs, and participants. This “orthogonal contrast” approach yielded *τ* = 0.12 seconds. At this *τ*, LSD was even more significantly reversible than placebo in all four conditions (*p*_Open_ < 0.0001, *p*_Closed_ < 0.0001, *p*_Music_ < 0.0001, *p*_Video_ = 0.0026) (**Supplementary Figure 4**).

The other approach employed cross-validation and permutation testing to select the *τ* that maximised the *difference* in irreversibility between placebo and LSD. (For most of the subsequent analyses, we used the *τ* that we derived from this approach. However, we doubt that we would have obtained significantly different results for the other analyses if we had opted for the orthogonal contrast approach, as irreversibility was consistently higher in placebo across all values of *τ*.) The cross-validation approach returned *τ* = 0.225 seconds. At this value of *τ*, the difference between LSD and placebo was actually smaller than it was when we used orthogonal contrast. Nevertheless, the decrease in irreversibility was highly significant across all conditions (*p*_Open_ < 0.0001, *p*_Closed_ < 0.0001, *p*_Music_ < 0.0001, *p*_Video_ = 0.0032) (**Figure 2a**; **Supplementary Table 1**).

**Figure 1.**
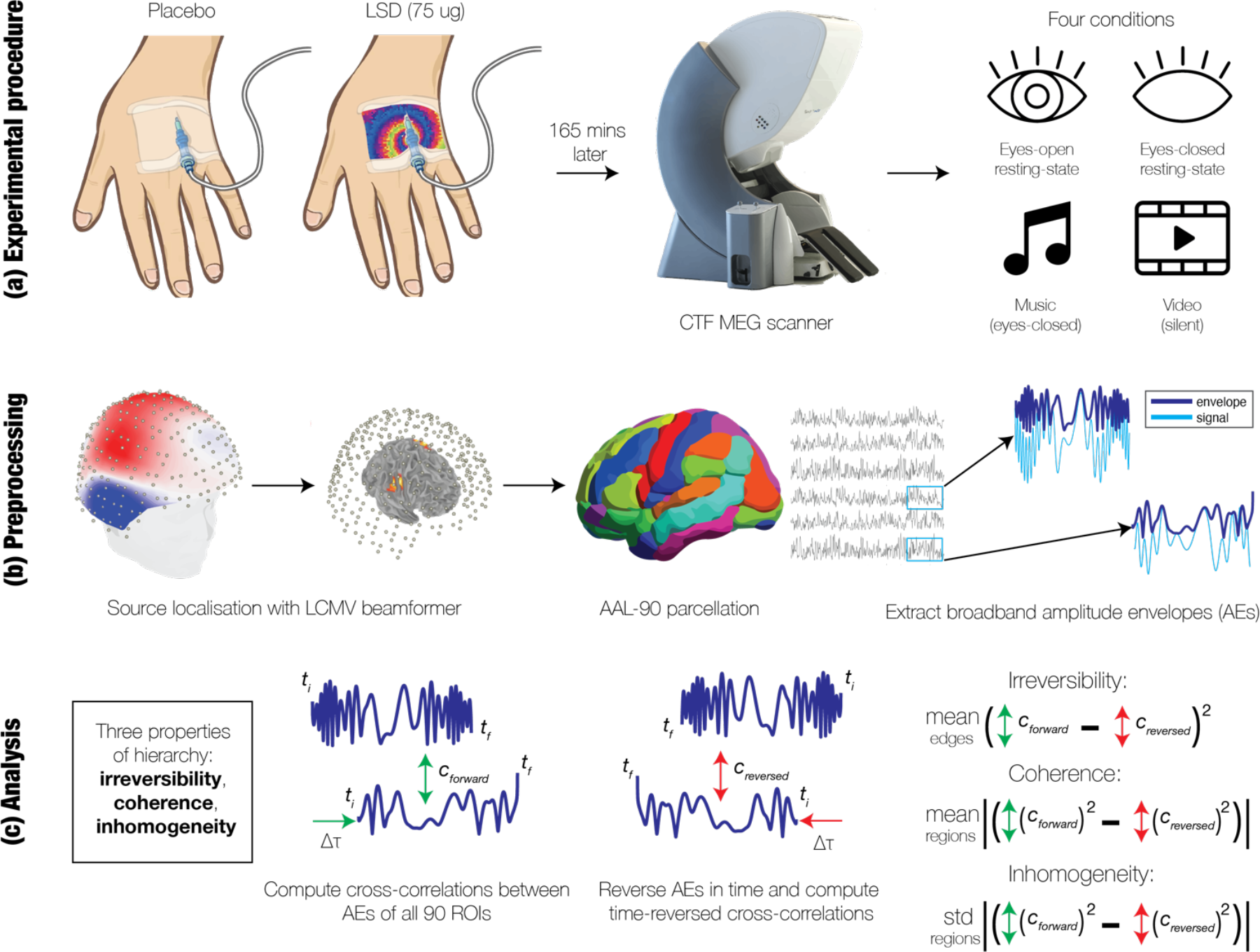
Overview. In this paper, we measure the effect of psychedelics on various measures of brain hierarchy. (a) Sixteen human participants were each administered placebo and 75 micrograms (ug) of LSD in two separate sessions (Carhart-Harris et al., 2016). Each session included four separate conditions: eyes-open resting-state, eyes-closed resting-state, listening to music with eyes closed, and watching a silent video. (b) The MEG data was preprocessed with methods described in a previous study (Mediano et al., 2024). Source localisation was performed using a linearly-constrained minimum variance (LCMV) beamformer. Broadband amplitude envelopes (AEs) were extracted via the Hilbert transform from 90 regional timeseries, which were parcellated with the Automated Anatomic Labelling (AAL) atlas. (c) We assessed three different properties of hierarchy: irreversibility, hierarchical coherence, and hierarchical inhomogeneity. We first measured time-delayed correlations, or cross-correlations, between the AEs of all 90 pairs of regions. We then reversed the AEs in time and performed the same time-delayed correlations. The irreversibility reflects the directedness of connectivity, i.e., the magnitude of the difference between connectivity from a region B to a region A and connectivity from A to B. Coherence and inhomogeneity are the mean and standard deviation of the incoming-outgoing asymmetry, i.e., the difference between the magnitude of incoming connections (B → A) and the magnitude of outgoing connections (A → B) at each region.

**Figure 2.**
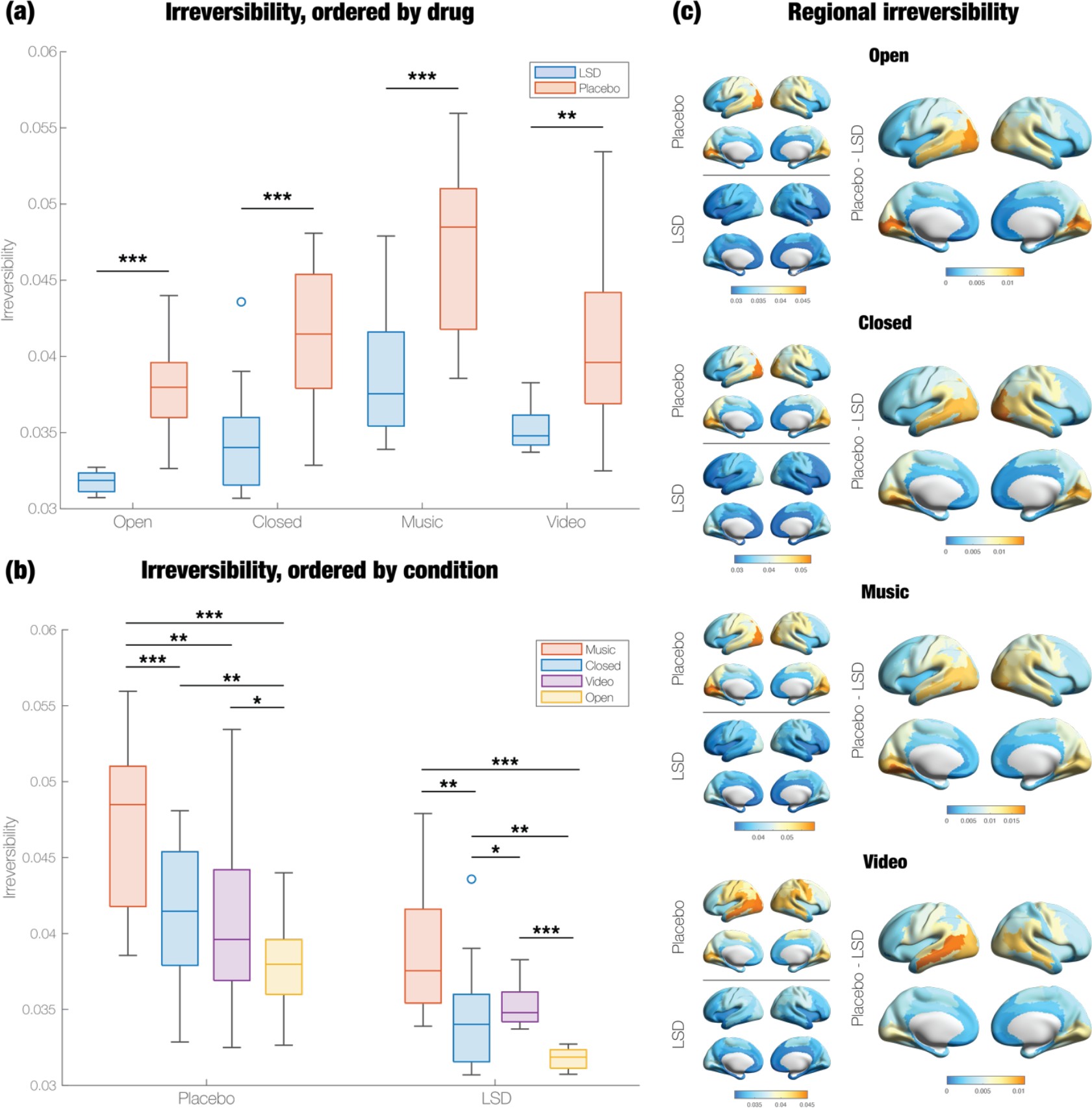
LSD makes broadband MEG signals significantly less directed (more reversible) across conditions. Using a cross-validation approach, we selected the value of the time-lag (τ = 0.225 seconds) that maximised the difference between irreversibility under LSD and irreversibility under placebo. We then computed the mean irreversibility across all pairs of regions at τ = 0.225 seconds. (a) MEG signals under LSD were significantly more reversible than under placebo, in all four conditions. (b) Under placebo, eyes-closed conditions (Music and Closed) are more irreversible than eyes-open conditions (Open and Video). (c) In addition to taking the mean across regions, we measured the irreversibility of each region. For each condition, we show the regional irreversibility under placebo and under LSD, as well as regions that display significant differences between placebo and LSD. Regional irreversibility follows an anteroposterior gradient; across conditions, it tends to be highest in occipital areas and lowest in frontal areas. Orange values correspond to larger differences and blue values to smaller differences. Most differences were significant, across conditions. *** p < 0.001, ** p < 0.01, * p < 0.05.

Under placebo, the conditions in which participants’ eyes were closed (Music and Closed) induced greater irreversibility than the conditions in which eyes were open (Video and Open) (**Figure 2b**). However, under LSD, Video was associated with significantly higher irreversibility than Closed.

In addition to taking the mean of irreversibility across all pairs of regions, we measured the irreversibility of each region. Regions in the occipital cortex consistently ranked the highest in irreversibility across all conditions, even those in which the participants’ eyes were closed (i.e., Closed and Music) (**Figure 2c**). In the Open, Closed, and Music conditions, all or nearly all (≥85) of the regions exhibit significantly reduced irreversibility on LSD compared to placebo. In the Video condition, 74 out of 90 regions had significantly less irreversible timeseries; the insignificant regions were primarily parietal. In none of the conditions were any of the regions significantly more irreversible on LSD compared to placebo. There is a robust anteroposterior gradient of changes in regional irreversibility, though it is more pronounced in the right hemisphere.

We also measured the frequency-specific effects of LSD on irreversibility. We band-pass filtered the MEG signal to four canonical frequency bands: theta (4-8 Hz), alpha (8-13 Hz), beta (13-30 Hz), gamma (30-48 Hz). We used cross-validation to determine the optimal *τ* for each frequency band. The alpha and beta results were consistent with those of our broadband analysis, but the theta and gamma bands were not (**Supplementary Figure 5**). LSD significantly increased irreversibility in all four of the conditions for both theta (*p*_Open_ = 0.0043, *p*_Closed_ = 0.0002, *p*_Music_ < 0.0001, *p*_Video_ = 0.0002) and in three conditions for gamma (*p*_Open_ = 0.0002, *p*_Closed_ < 0.0001, *p*_Music_ = 0.0006, *p*_Video_ = 0.2078). However, in the theta band, LSD is more irreversible than placebo for values of *τ* ranging from 0.005 seconds to at least 0.25 seconds. Then, placebo is more irreversible than LSD until *τ* is at least 0.625 seconds. Since 0.25 seconds is the length of one 4Hz theta cycle, these results may indicate that there is greater asymmetry in within-cycle connectivity on LSD and larger asymmetry in between-cycle connectivity on placebo.

One problem with time-delayed correlations is that they do not control for confounding influences. There could be a strong correlation between a region *A* and a region *B* even if there were no causal relationship between the two, so long as there is a region *C* that is driving activity in both. Partial time-delayed correlations between any two timeseries provide a solution to this limitation by regressing out all other regions. With this approach, we still found that LSD significantly reduces irreversibility across all four conditions (*p*_Open_ < 0.0001, *p*_Closed_ < 0.0001, *p*_Music_ < 0.0001, *p*_Video_ = 0.004) (**Supplementary Figure 6**).

### Section 3.2. LSD decreases hierarchical coherence and inhomogeneity

Irreversibility captures the *connections* that exhibit the largest differences between their incoming component and outgoing component. Here, we measured the difference in magnitude between the strongest incoming and outgoing connections *at each region*. While this incoming-outgoing asymmetry appears to be similar to irreversibility, **Figure 3a** illustrates toy models in which these two quantities diverge from each other. In particular, the direction of connectivity between the top right and top left node changes between network (1) and (2), such that the top right node only has incoming connections in network (2); however, the irreversibility of all the connections remains the same in both networks.

**Figure 3.**
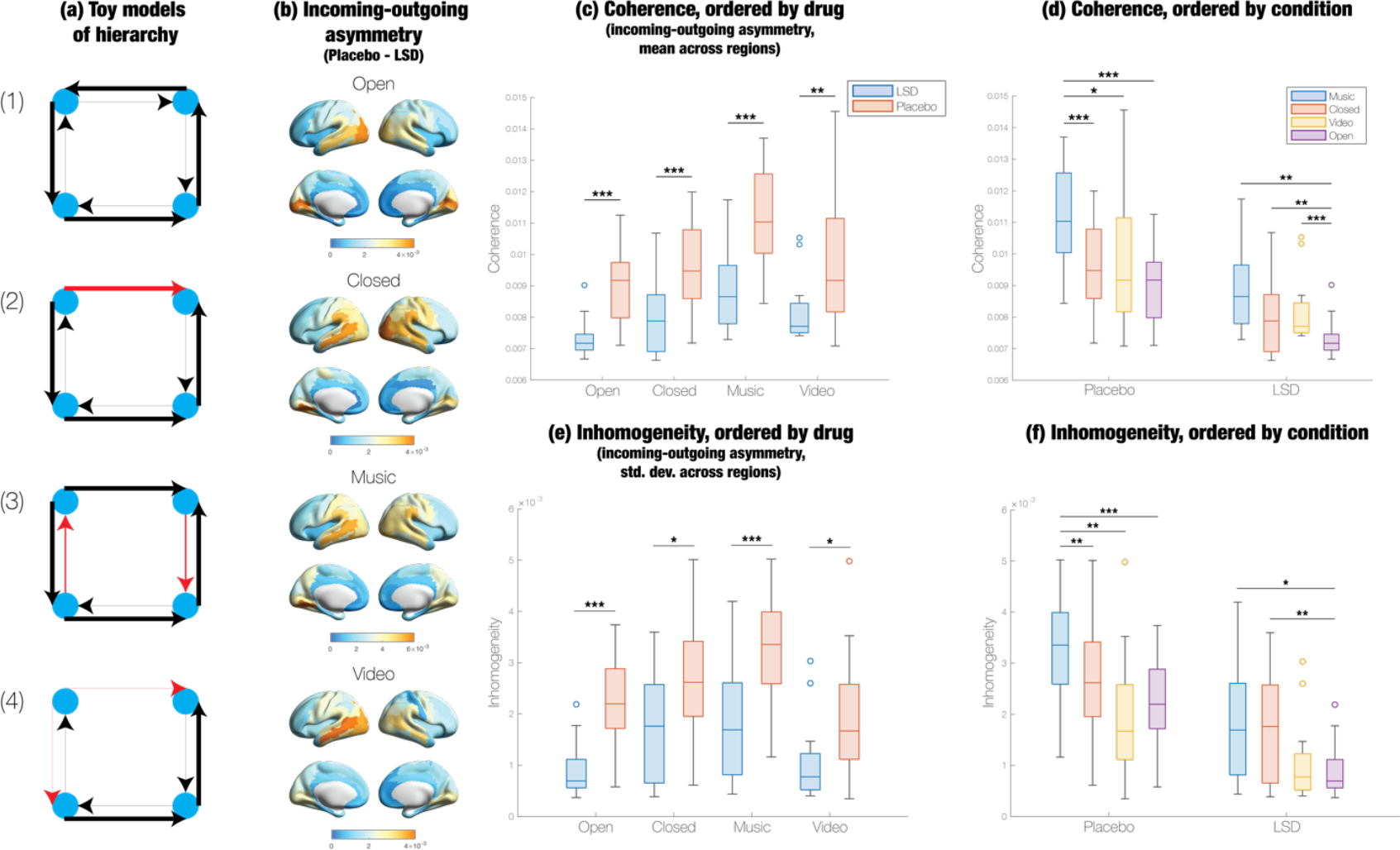
LSD significantly reduces hierarchical coherence and inhomogeneity across conditions. (a) Despite being irreversible, toy model (1) is completely non-hierarchical because each node is receiving the same amount of information that it is sending; the total weights of the incoming and outgoing connections are equally strong for all nodes. By altering the direction of a single edge (shown in red), model (2) is more hierarchical because the incoming and outgoing connections are now imbalanced for the upper left and upper right nodes. Its hierarchical coherence and inhomogeneity – mean and standard deviation of the incoming-outgoing asymmetry, respectively – are higher than in model (1). Model (3) has higher hierarchical coherence but lower hierarchical inhomogeneity than (2) because the red edges grow larger in magnitude. Conversely, model (4) has lower hierarchical coherence but higher hierarchical inhomogeneity than (3) because the weights of the red edges decrease. Both coherence and inhomogeneity are required for hierarchy; therefore, both models (3) and (4) are less hierarchical than model (2). (b) The incoming-outgoing asymmetry is significantly higher under placebo in most regions; furthermore, the spatial topography of the incoming-outgoing asymmetry is very similar to that of regional irreversibility. (c) In all four conditions, LSD significantly reduces hierarchical coherence. (d) Under LSD but not placebo, hierarchical coherence is significantly higher in the Closed condition than in the Open condition. (e) Hierarchical inhomogeneity significantly decreases under LSD in all four conditions. (f) As with hierarchical coherence, inhomogeneity is significantly greater in Closed compared to Open, but only under LSD. *** p < 0.001, ** p < 0.01, * p < 0.05.

**Figure 3b** displays significant differences in the incoming-outgoing asymmetry between placebo and LSD. LSD significantly decreases the asymmetry across all regions, under all four experimental conditions. The spatial distribution of these asymmetry values is nearly identical to that of regional irreversibility; the asymmetry tends to be highest in occipital and temporal cortices and lowest in frontal cortices. Thus, incoming-outgoing asymmetry and irreversibility typically do not diverge from each other.

We define hierarchical coherence and inhomogeneity as the mean and standard deviation of the incoming-outgoing asymmetry across regions, respectively. As shown in **Figure 3c-f** and **Supplementary Tables 2-3**, LSD significantly reduces both hierarchical coherence (*p*_Open_ < 0.0001, *p* _Closed_ = 0.0002, *p* _Music_ < 0.0001, *p* _Video_ = 0.0062) and hierarchical inhomogeneity (*p* _Open_ < 0.0001, *p*_Closed_ = 0.0139, *p*_Music_ = 0.0003, *p*_Video_ = 0.0121) in all four conditions. Under both placebo and LSD, brain activity was more coherent and inhomogeneous in the eyes-closed conditions (Music and Closed) than the eyes-open conditions (Video and Open), though the difference between Closed and Open for both hierarchy metrics was significant only on LSD.

### Section 3.3. The measures of hierarchy dynamically reconstitute themselves over time

The above analyses were static, so we performed a dynamical analysis to determine how the three hierarchy metrics evolve over time. First, we computed each metric in overlapping sliding windows of 1-second length. **Supplementary Figure 7a** shows the irreversibility timeseries in one condition (Open) for a particular participant, under both placebo and LSD. To inspect the dynamics, we performed a time-delay embedding of the irreversibility timeseries, which is a common procedure for extracting latent dynamical variables (Takens, 1981; Varley & Sporns, 2021). Each embedding dimension *d* represents the metric at time (*d* – 1) * *τ*_RN_ (**Supplementary Figure 7b**). (Note that *τ*_RN_ is distinct from the *τ* determined in *Section 3.1*.) It is clear that points in the time-delay embedding are closer together under LSD than under placebo, suggesting that the dynamics of irreversibility are more similar on LSD compared to placebo. To quantify the similarity of the dynamics, we constructed a recurrence network, in which timepoints in the centrality timeseries are connected by an edge if their distance in the time-delay embedding is below a certain threshold. In the recurrence networks displayed in **Supplementary Figure 7c**, the density of edges, or recurrence rate, is evidently much higher under LSD for this particular participant. When we consider all participants, the recurrence rate of the irreversibility dynamics is significantly higher on LSD in all four conditions (*p*_Open_ = 0.0001, *p*_Closed_ = 0.0185, *p*_Music_ = 0.0032, *p*_Video_ = 0.0416) (**Supplementary Figure 7d**). When we repeat the above steps for the other two hierarchy metrics, we find that LSD significantly increases the recurrence rate under all conditions (hierarchical coherence: *p*_Open_ = 0.0013, *p*_Closed_ = 0.0002, *p*_Music_ < 0.0001, *p*_Video_ = 0.0352; hierarchical inhomogeneity: *p*_Open_ < 0.0001, *p*_Closed_ = 0.0030, *p*_Music_ = 0.0007, *p*_Video_ = 0.0012) (**Supplementary Figure 7e-f**).

### Section 3.4. Irreversibility distinguishes LSD and placebo more than other measures of functional connectivity and hierarchy

How do the three hierarchy metrics compare to other measures of functional connectivity (FC), such as undirected FC and cross-correlations? To answer this question, we first measured the effects of LSD on the latter two measures. Over all conditions, LSD significantly reduced each metric, both when averaged across regions and within all or most regions (**Supplementary Figure 8**). Furthermore, the spatial topographies of regional changes in undirected FC and forward cross-correlations were strikingly similar to those of regional irreversibility. They were lowest in frontal cortices and highest in occipital and temporal regions, though decreases in undirected FC were more pronounced in occipital cortices, compared to irreversibility.

Yet despite the similarity between the topographies of irreversibility, forward cross-correlations, and undirected FC, mean irreversibility differentiates LSD and placebo much better than the mean of the other two metrics. We trained random forest classifiers to distinguish placebo from LSD data based on regional irreversibility or one of these other metrics of FC. The classifier trained on irreversibility significantly outperformed the classifiers trained on forward cross-correlations and undirected FC, in addition to those trained on the other two hierarchy metrics (**Figure 4**).

**Figure 4.**
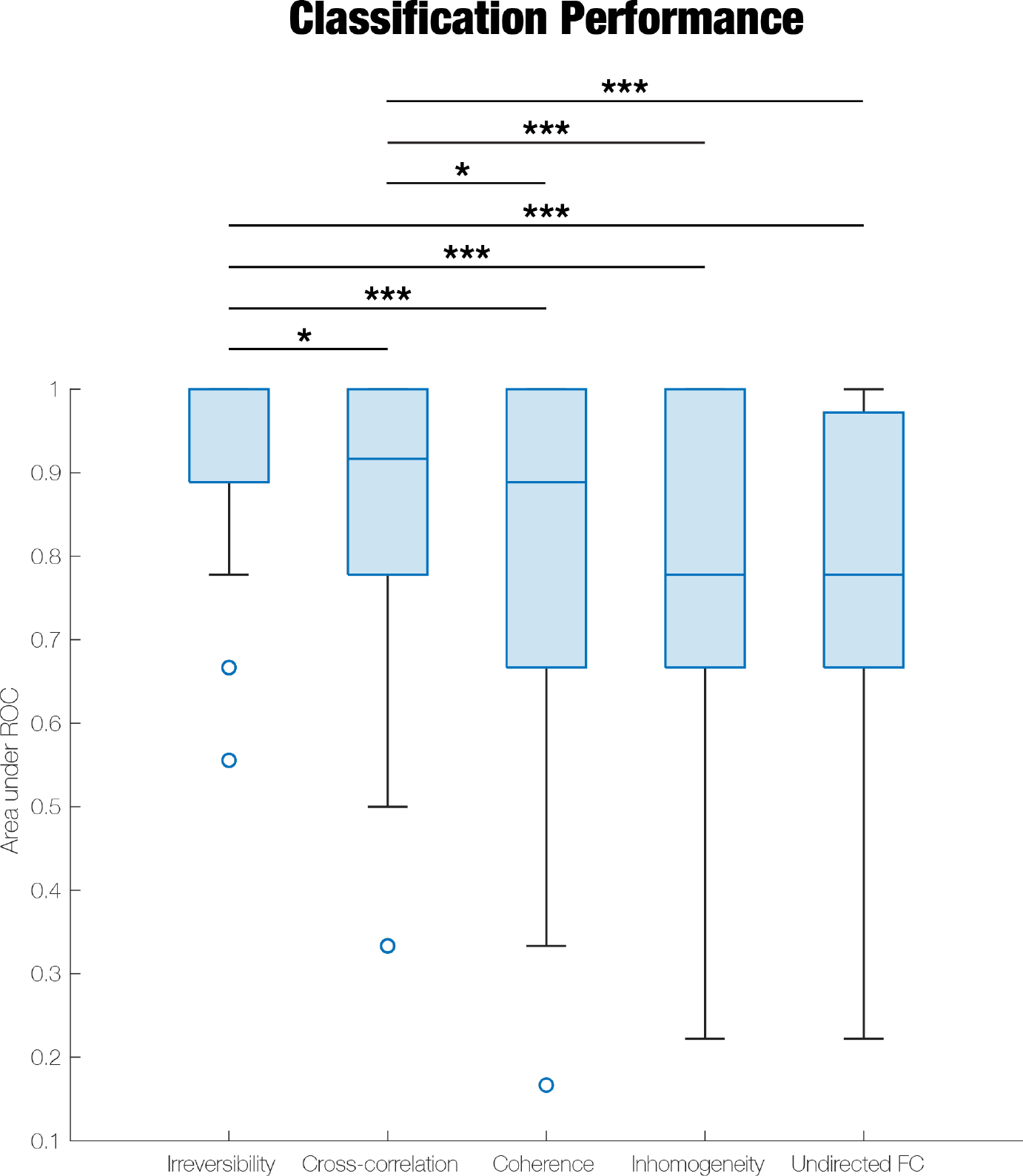
Regional irreversibility distinguishes LSD and placebo better than other metrics of functional connectivity. We quantified the effect of LSD on two different metrics of functional connectivity (FC): undirected FC (Pearson correlation) and cross-correlations (measured only in the forward direction of time). The results of this analysis are shown in **Supplementary Figure 8**. We then trained a random forest classifier to distinguish between LSD and placebo on the basis of undirected FC, cross-correlations, and the three hierarchy metrics introduced in this paper. Classification accuracy, as measured by the area under the Receiver Operating Characteristic (ROC) curve, was significantly higher for the classifier trained on irreversibility, compared to all other classifiers. *** p < 0.001, ** p < 0.01, * p < 0.05.

### Section 3.5. Regional irreversibility does not correlate with serotonin receptor expression

Because the hallucinogenic effects of psychedelics are attributed to their interactions with the 5-HT_2A_ receptor (Vollenweider et al., 1998), we hypothesised that the spatial patterns of changes in regional irreversibility would be correlated with 5-HT_2A_ receptor expression. However, this was not the case in any of the conditions (*p*_Open_ = 0.9560, *p*_Closed_ = 0.9020, *p*_Music_ = 0.9560, *p*_Video_ = 0.8100) (**Supplementary Figure 9a-b**).

We also anticipated that the difference in regional irreversibility between placebo and LSD would be correlated more with 5-HT_2A_ receptor expression than with 5-HT_1A_ or 5-HT_1B_ expression, as these receptors do not seem to play as essential of a role in mediating the hallucinogenic effects of psychedelics in humans (Nichols, 2016). However, we found no significant correlations (5-HT_1A_: *p*_Open_ = 0.2130, *p*_Closed_ = 0.2150, *p*_Music_ = 0.2250, *p*_Video_ = 0.2500; 5-HT_1B_: *p*_Open_ = 0.3360, *p*_Closed_ = 0.3180, *p*_Music_ = 0.3340, *p*_Video_ = 0.4360) (**Supplementary Figure 9c-f**).

## Section 4. Discussion

Our main finding is that LSD makes broadband neural signals more reversible across different experimental conditions: eyes-open resting-state, eyes-closed resting-state, listening to music, and watching a silent video. We also found that LSD significantly decreases two other properties of hierarchy: hierarchical coherence and inhomogeneity, or the mean and standard deviation of the asymmetry between incoming and outgoing connections. While the spatial topography of changes in irreversibility is similar to that of other FC metrics, such as undirected FC and cross-correlations, we found that random forest algorithms classify placebo and LSD more accurately when trained on irreversibility.

### Section 4.1. Psychedelics and hierarchy

Here, we aimed to produce a simple method for testing the hypothesis that psychedelics flatten the hierarchy of the brain (R. L. Carhart-Harris & Friston, 2019). Our method is predicated on the assumption that any hierarchy must exhibit asymmetries in directed information flow; that is, higher-order components control the activity of lower-order components, but not vice versa. Based on this assumption, we investigated three properties of hierarchy: irreversibility, hierarchical coherence, and hierarchical inhomogeneity.

Our finding that LSD significantly reduces directional asymmetry, or irreversibility, is consistent with previous MEG studies, which showed that LSD and psilocybin reduce other measures of directed connectivity like Granger causality across the whole brain (Barnett et al., 2020). Other work has also demonstrated that the psychedelic DMT amplifies bottom-up travelling waves while reducing the power of top-down travelling waves. While there have been an abundance of studies on undirected FC under psychedelics (Barrett, Doss, et al., 2020; Barrett, Krimmel, et al., 2020; Bedford et al., 2023; R. L. Carhart-Harris et al., 2016; Dai et al., 2023; Gaddis et al., 2022; Grimm et al., 2018; Madsen et al., 2021; Mason et al., 2020; Palhano-Fontes et al., 2015; Pallavicini et al., 2019; Roseman et al., 2014; Timmermann et al., 2023), there have been relatively few studies on directed FC by comparison (Alamia et al., 2020; Barnett et al., 2020; de Araujo et al., 2011; Timmermann et al., 2023). However, the fact that irreversibility distinguishes placebo and LSD better than undirected measures, according to our random forest analyses, suggests that irreversibility may be a more specific biomarker of psychedelics. It is worth noting that a handful of studies have measured directed, *effective* connectivity with dynamic causal modelling (DCM), but the examined regions of interest differ widely between studies (Bedford et al., 2023; Kaelen et al., 2016; Kraehenmann et al., 2015; Preller et al., 2019; Timmermann et al., 2018). Nevertheless, all the studies demonstrated that psychedelics have directionally asymmetric effects on connectivity.

However, irreversibility does not always imply hierarchy. For instance, in network (1) of the toy models shown in **Figure 3a**, all connections are highly irreversible, yet the network is not hierarchical at all, since information flows towards and away from all nodes equally. But network (2) is more hierarchical because one of the connections (shown in red) flips direction, such that the top right node receives more information than it transmits, and vice versa for the top left node. Network (2) has higher hierarchical coherence and inhomogeneity than network (1) because there is a non-zero asymmetry between incoming and outgoing connections at these two nodes. Furthermore, non-zero coherence or inhomogeneity always entails irreversibility since the former implies a pairwise asymmetry in connectivity. We propose that both hierarchical coherence and inhomogeneity are necessary for hierarchy. For example, network (3) exhibits increased hierarchical coherence but decreased inhomogeneity relative to network (2), and network (4) higher inhomogeneity but lower coherence compared to network (2). Yet both are evidently less hierarchical than network (2). In network (2), the top left node is predominantly a transmitter of information, whereas the top right is predominantly a receiver. However, the top left node receives more and the top right node transmits more in network (3), whereas the top left node transmits less and the top right node receives less in network (4). The significant decreases in not only irreversibility but also hierarchical coherence and inhomogeneity under LSD lend strong support to the claim that this psychedelic diminishes hierarchy.

While these toy models illustrate instances in which irreversibility and the incoming-outgoing asymmetry dissociate from each other, they exhibit very similar spatial patterns in empirical data. In other words, although it is theoretically possible for brain regions to become more irreversible but also more balanced in its incoming and outgoing connections, this does not appear to occur in the data at hand.

Our assumption that hierarchies exhibit asymmetry between incoming and outgoing connectivity is equivalent to the statement that hierarchies must contain acyclic information flow, a definition that is made explicit in measures of flow hierarchy (Luo & Magee, 2011). Here, the strength of hierarchy corresponds to the proportion of edges that do not participate in cycles.

Another established measure of hierarchy defines hierarchical coherence in a similar way. So-called “trophic coherence” is the normalised difference between “trophic levels,” or hierarchical levels, which are essentially determined based on both the incoming-outgoing asymmetry and the irreversibility (MacKay et al., 2020). However, trophic coherence can only be measured on a non-negative network, whereas the time-delayed correlations that we measured in this study can be negative. In the future, we hope to use a whole-brain modelling approach to extract an effective, non-negative network from the same dataset, so that we can subsequently quantify trophic coherence (Jobst et al., 2017; Kringelbach et al., 2023; Kringelbach & Deco, 2020).

A recent graph-theoretic framework that generalises trophic coherence bears even greater resemblance to our definition of hierarchy (Moutsinas et al., 2021). Here, the “democracy coefficient” of a network is defined based on the mean of the incoming and outgoing connections between hierarchical levels, and the “hierarchical incoherence” on the standard deviation of the same connections. Similar to trophic coherence, hierarchical levels are identified by solving a linear equation that essentially relates irreversibility to the in- and out-degrees of the network. Like ours, this framework acknowledges that both the mean and the standard deviation of imbalance between incoming and outgoing connections characterise the strength of a hierarchy.

So far, the notions of hierarchy that we have considered are statistical or functional rather than anatomical. Anatomical hierarchies do not always align with functional hierarchies (Silvanto, 2015; Zeki, 2018). However, the rank-ordering of irreversibility is congruent with the anatomical hierarchy of cortical visual regions, in which the primary visual cortex (V1) is at the bottom and areas connected to the dorsal and ventral visual processing streams (e.g., the frontal eye field [FEF]) are towards the top (Felleman & Van Essen, 1991). In the Open, Closed, and Music conditions, the calcarine, which contains V1 (Galetta, 2014), is either the most or the second most irreversible region. The cuneus and lingual gyrus, which encompass V2, V3, and V4 (Palejwala et al., 2021), are more reversible, followed by the precentral gyrus, which contains a portion of the FEF (El-Baba & Schury, 2024). In the Video condition, the middle temporal gyrus, which contains V5, is more irreversible than the calcarine; this is consistent with the fact that V5 controls motion perception (Zeki, 2015).

The direction of the hierarchy defined in this study appears to contradict the traditional direction of visual hierarchies; here, V1 is near the top, whereas V1 is usually at the bottom. Our analysis is agnostic about whether regions that transmit or receive information are higher or lower in the hierarchy; rather, it places regions with a high incoming-outgoing asymmetry at the top of the hierarchy. It turns out that the brain areas with the largest asymmetry, i.e., visual areas, transmit more information than they receive. Our finding that outgoing connectivity is highest in visual areas is consistent with previous research on directed connectivity (Deco et al., 2021).

Finally, we also show that the brain “explores” a broader set of hierarchical dynamics under placebo than LSD. While previous studies showed that LSD expands the repertoire of connectivity motifs (Atasoy et al., 2017; R. Carhart-Harris et al., 2014; Tagliazucchi et al., 2014), they averaged connectivity over time and therefore did not consider dynamics. Studies that investigated dynamics either found no differences between psychedelics and placebo or did not explicitly measure the dynamics of hierarchy (Luppi et al., 2021; Timmermann et al., 2023). To our knowledge, this is the first study that assessed the effect of psychedelics on the dynamics of hierarchy. Our finding suggests that hierarchy is not static but rather changes over time, yet LSD constrains the evolution of the hierarchy to a smaller subset of dynamics.

### Section 4.2. Entropy, entropy production, and the arrow of time

Ever since Robin Carhart-Harris formulated the Entropic Brain Hypothesis, which claims that the altered state of consciousness on psychedelics is encoded by a brain state of elevated entropy (Carhart-Harris et al., 2014), a number of fMRI, MEG, and EEG studies have investigated the effect of psychedelics on entropy (Alonso et al., 2015; Lebedev et al., 2015, 2016; Mediano et al., 2024; Ruffini et al., 2023; Schartner et al., 2017; Singleton et al., 2022; Tagliazucchi et al., 2014; Timmermann et al., 2019, 2023; Varley et al., 2020; Viol et al., 2017, 2019, 2023). Virtually all of them demonstrated that psychedelics increase entropy, though they do not use consistent techniques and are mostly secondary analyses of the same two datasets (R. L. Carhart-Harris et al., 2012, 2016). (In fact, a study on a new dataset found that psilocybin did not have a significant effect on seven of the 12 entropy measures in the literature (McCulloch et al., 2023)).

However, to our knowledge, none of these studies have measured the effect of psychedelics on entropy *production*. The Crooks fluctuation theorem in physics states that entropy production is exponentially related to irreversibility, i.e. the ratio between the forward and time-reversed transition probabilities, when the dynamics of the system are Markovian (the state at each timepoint only depends on the state at the previous timepoint) (Crooks, 1998).

It is important to note that entropy and entropy production are not equivalent. Entropy refers to the variability or unpredictability of states of a system. It is highest at thermodynamic equilibrium, which is a state of maximum disorder; the system is capable of existing in a maximally large number of states. For example, once a drop of ink has diffused in a tank of water and reached equilibrium, it has evenly spread out and has therefore maximised its probability of being located at any position in the tank. Due to the Second Law of Thermodynamics, entropy (in a closed system) always increases over time; the system, e.g., the drop of ink, tends toward thermodynamic equilibrium. Thus, once entropy has reached its maximum in equilibrium, it cannot do anything other than remain constant. Entropy is maximal while entropy production is minimal (zero). Equilibrium also respects detailed balance; every forward transition between states is balanced by the corresponding reverse transition (Lewis, 1925). Exiting equilibrium implies that detailed balance is broken and therefore that the dynamics of the system become irreversible.

Outside of equilibrium, entropy and entropy production may also change in opposite directions. For instance, MEG and EEG studies have found that psychedelics increase Lempel-Ziv complexity (LZc) (Mediano et al., 2024; Schartner et al., 2017; Timmermann et al., 2019), which is known to be an efficient estimator of entropy rate, or the amount of uncertainty about the value of the next sample in a data timeseries (Mediano et al., 2023; Ziv & Lempel, 1978). LZc should be higher in reversible systems, in which the state that the system will transition to at the next timepoint is maximally uncertain. (For instance, in a cyclic and reversible system, it is equally probable that information will flow clockwise or counterclockwise at the next timepoint.) Intriguingly, according to a recent analysis of the same primary dataset that we examined, LZc is higher in eyes-open conditions than eyes-closed conditions, under both placebo and LSD (Mediano et al., 2024). On the other hand, we demonstrated that irreversibility is higher in eyes-closed conditions than eyes-open conditions. This is consistent with the claim that LZc and irreversibility are inversely related.

Our measure of entropy production, INSIDEOUT, does not quantify the probability of forward and backward transitions between brain *states*; instead, INSIDEOUT only computes the forward and backward time-delayed correlations between brain *regions*. In the future, we aim to use the Ornstein-Uhlenbeck (OU) model, which analytically derives entropy production and has recently been applied to fMRI timeseries (Gilson et al., 2023), to directly obtain the amount of entropy production in the same dataset. Furthermore, recent work has applied the OU model to demonstrate that hierarchical network structure drives entropy production (Nartallo-Kaluarachchi et al., 2024).

### Section 4.3. Limitations

While we have shown here that psychedelic use is associated with directedness of brain connectivity, we have not yet demonstrated causality. REBUS proposes that the flattened hierarchy gives rise to both the altered states of consciousness and therapeutic effects of psychedelics, but we have not provided evidence that the subjective experience of psychedelics is governed by the three hierarchical metrics that we investigated.

Additionally, we measured the three properties of hierarchy in all brain regions, yet a smaller subset of latent features or brain regions likely mediates directed connectivity (Cai et al., 2021; Taghia et al., 2018; Vidaurre et al., 2018). While we averaged across the top 5% of values in each data epoch to compute the hierarchical properties, we would ideally apply a more principled method for reducing dimensionality and noise, such as reduced rank regression (MacDowell et al., 2023).

Finally, increases in hierarchical coherence and inhomogeneity may not always correspond to increases in the strength of hierarchy. For instance, consider a network with horizontal (within-layer) and vertical (between-layer) connections. The hierarchical flow of information is vertical in this network; information tends to flow upwards towards a small set of higher-order nodes. According to our framework, perturbing the imbalance of a node by altering its horizontal connectivity would have the same effect on the hierarchy as changing its vertical connectivity would. However, in reality, only the vertical connectivity modulates the strength of hierarchy in this network.

## Section 5. Conclusion

Several studies have examined the effects of psychedelics on entropy, but none have analysed their effects on entropy *production*. Whereas entropy reflects the amount of variability or unpredictability in a system, entropy production is the amount of irreversibility in the system’s dynamics. Recent work has shown that hierarchical network structure drives entropy production by increasing the directional asymmetry of network connectivity (Nartallo-Kaluarachchi et al., 2024). Here, we demonstrate that LSD reduces the irreversibility, or directional asymmetry, of brain activity, thereby decreasing entropy production. We also link irreversibility and the imbalance between incoming and outgoing connectivity to the strength of functional hierarchy. Thus, our results corroborate the hypothesis that psychedelics flatten hierarchy and reduce the imbalance between top-down and bottom-up information flow.

