## Supplementary Materials for "LSD flattens the hierarchy of directed information flow in fast whole-brain dynamics"

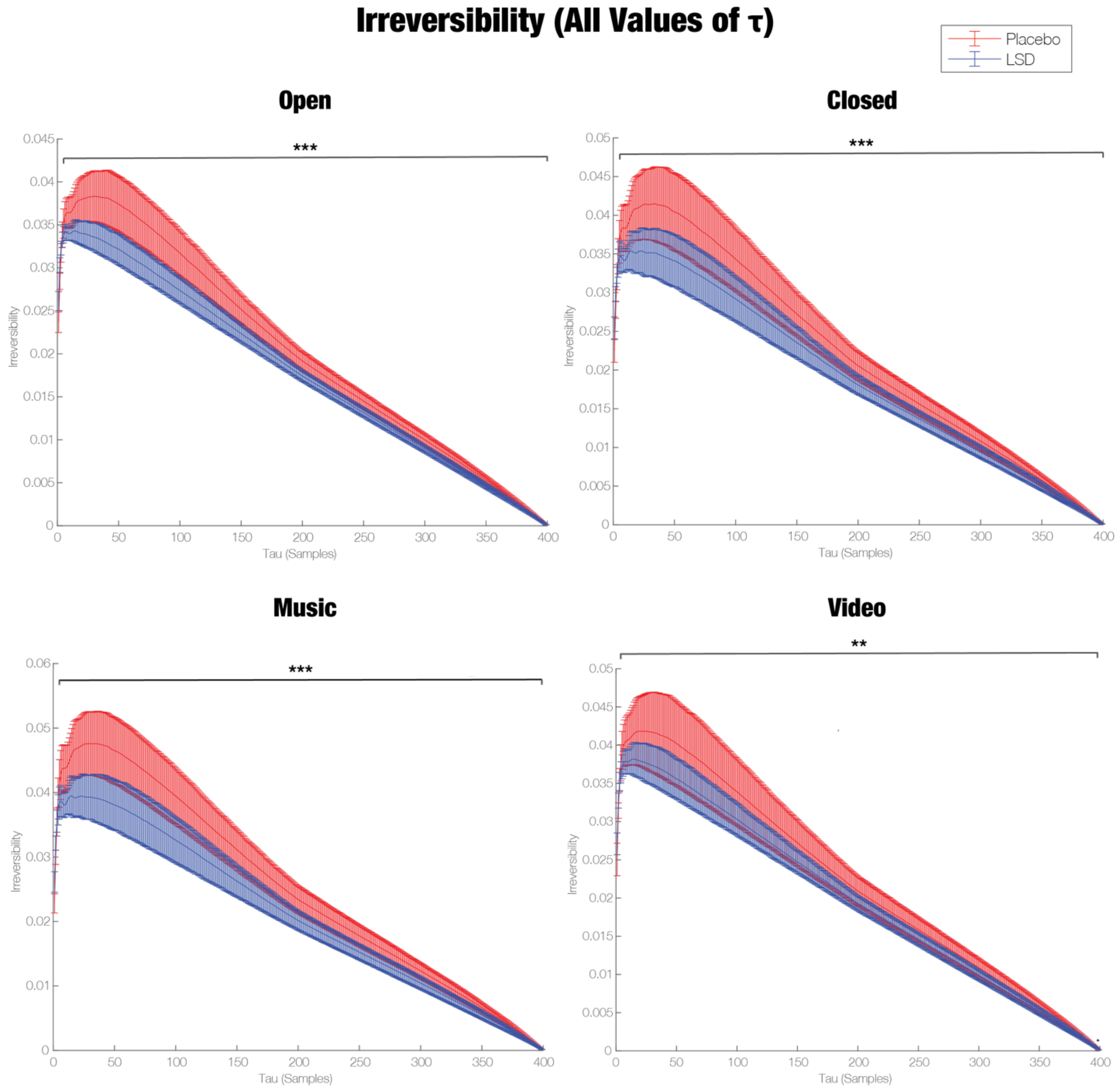


***Supplementary Figure 1. LSD reduces irreversibility across all values of τ**. Irreversibility is calculated as mean of the top 5% of squared differences between forwards and time-reversed cross-correlations of regional MEG amplitude envelopes. Cross-correlations are correlations shifted by an interval τ. We first computed irreversibility across all possible values of τ. (Since we executed INSIDEOUT on each epoch of data, which consisted of 400 samples (2 seconds), there were 399 possible values of τ.) We used cluster-based permutation testing to identify the values of τ that were associated with significant differences in irreversibility between LSD and placebo. Across nearly all values of τ, LSD was significantly more reversible than placebo in all four of the conditions. LSD was insignificantly more irreversible than placebo only at values of τ ranging from 1-3 samples (0.005-0.015 seconds). *** p < 0.001, ** p < 0.01, * p < 0.05.*

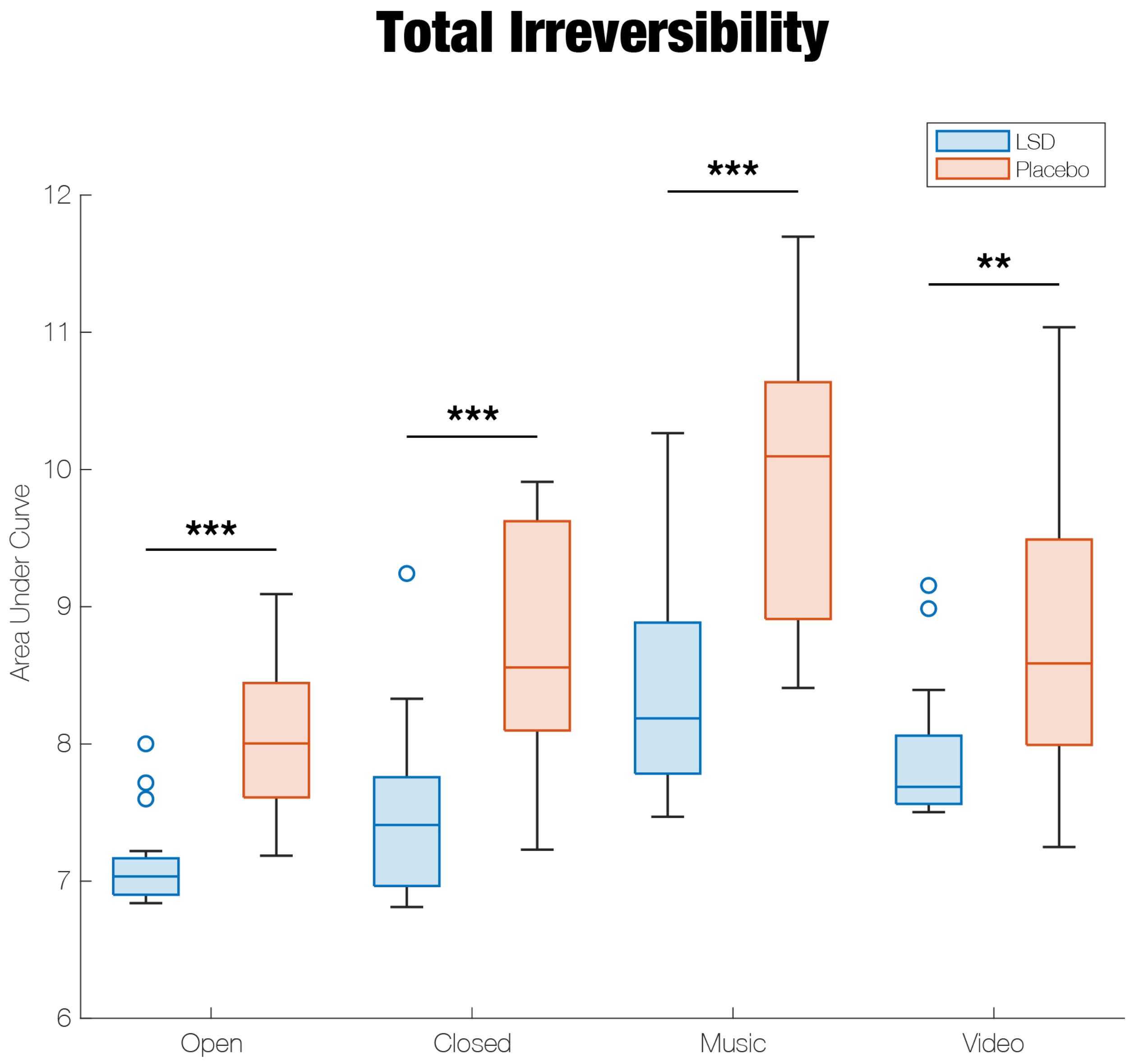


***Supplementary Figure 2. LSD decreases the "total" amount of irreversibility across all τ.*** *We integrated the value of irreversibility across all possible values of τ (0.005-1.995 seconds) by computing the area under the τ vs irreversibility curve (**Supplementary Figure 1**). In line with our main result, LSD is significantly more reversible than placebo across all four conditions.* *** $p < 0.001$, ** $p < 0.01$, * $p < 0.05$.

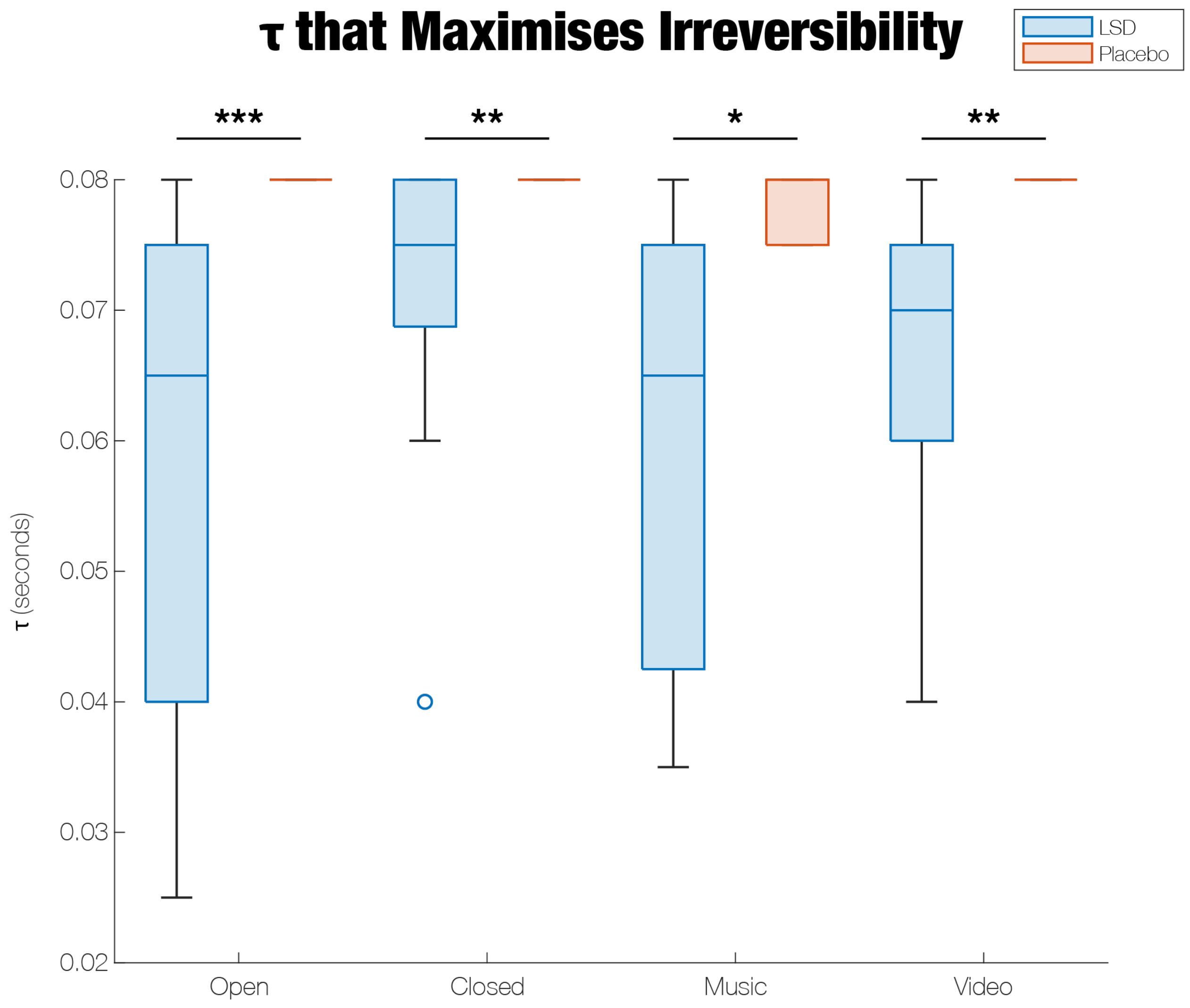


***Supplementary Figure 3. LSD increases the spread of the values of τ that maximise irreversibility.*** *As shown in **Supplementary Figure 1**, irreversibility peaks at a single value of τ. In the Open, Closed, and Video conditions, the τ that yielded the peak value of irreversibility was the same across all participants under placebo (τ = 0.12 seconds). However, under LSD, participants exhibited a significantly lower and much wider range of τ values that maximised irreversibility. If τ reflects the length of time that it takes regions to propagate signals to one another, these results perhaps suggest that there is greater inter-subject variability in the speed of connectivity under LSD. An alternative explanation is that connectivity becomes more local for some participants than others, thereby shortening their τ by a greater amount.* *** $p < 0.001$, ** $p < 0.01$, * $p < 0.05$.

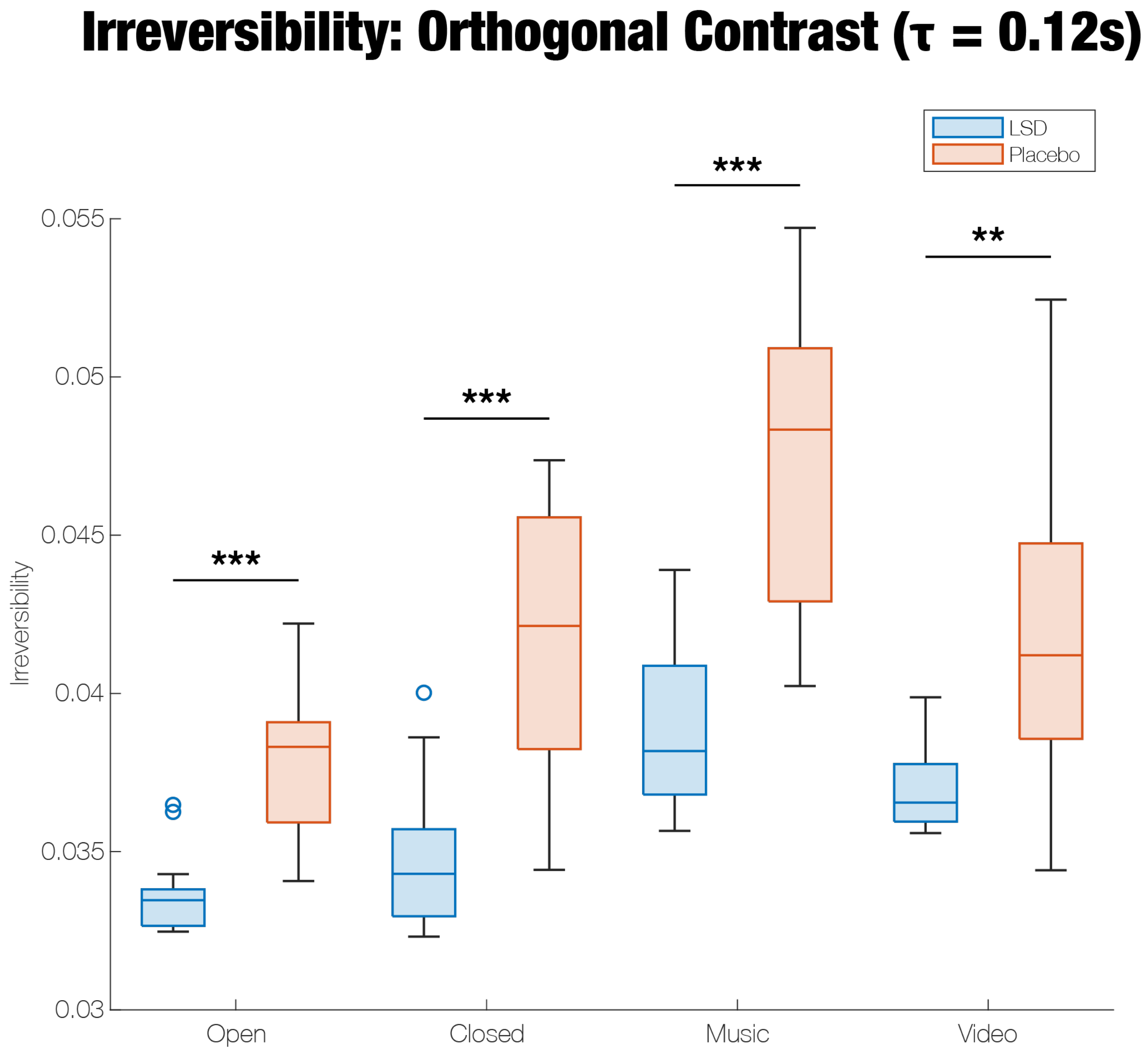


***Supplementary Figure 4. Using a different approach for selecting τ, we still find that the LSD data is significantly more reversible than the placebo data.*** *We used two different methods to identify a specific τ for conducting subsequent analyses. (Using a single τ expedited the analyses.) The results from the first method, which uses cross-validation to find the τ that maximally distinguishes LSD and placebo, are described in the main text. The second method, which we refer to as the orthogonal contrast technique, identifies the τ that maximises the mean irreversibility across all datasets. The cross-validation approach yielded τ = 0.225 seconds, whereas the orthogonal contrast approach yielded τ = 0.12 seconds. The results achieved with τ = 0.12 were very similar to those that we obtained with τ = 0.225. LSD was significantly more reversible across all four conditions. *** p < 0.001, ** p < 0.01, * p < 0.05.*

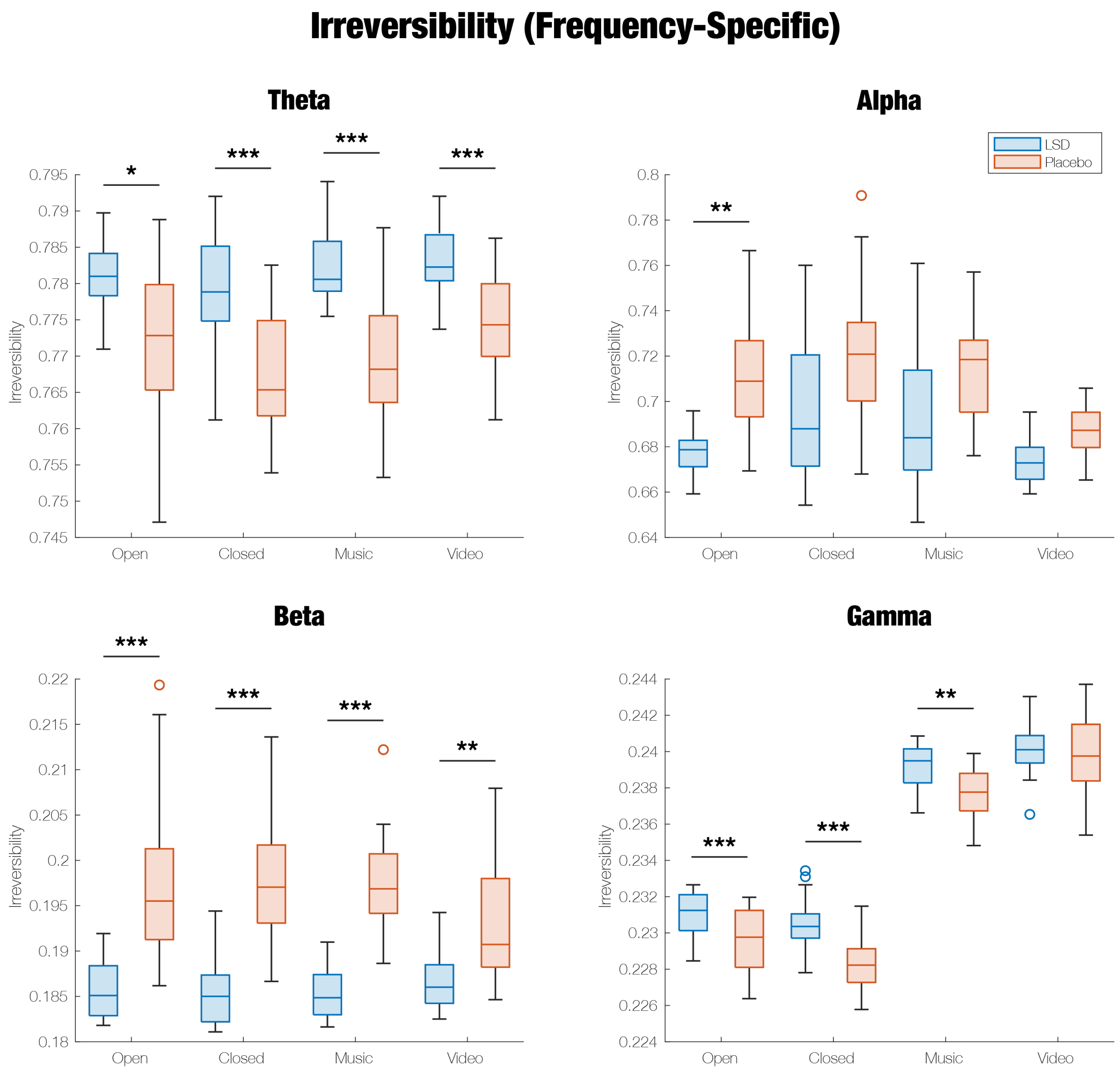


***Supplementary Figure 5. LSD is significantly more reversible than placebo in the alpha and beta bands, but not in the theta and gamma bands.*** *After obtaining results for the broadband data, we bandpass-filtered the MEG timeseries into four frequency bands: theta (4-8 Hz), alpha (8-13 Hz), beta (13-30 Hz), and gamma (30-48 Hz). (We chose not to analyse the delta band because each epoch of data in each dataset was only two seconds long, which is equivalent to one or two low-frequency delta cycles.) In alpha and beta, LSD was more reversible than placebo, although this effect was only significant in the eyes-open condition in the alpha band. However, the theta and gamma results showed the opposite effect. That being said, in the theta band, LSD only ranked higher than placebo in irreversibility at certain values of* $\tau$*. In particular, the irreversibility of LSD was greater than that of placebo for* $\tau$ *under 0.25 seconds, but smaller for* $\tau$ *between 0.25 and 0.625 seconds. Since 0.25 seconds is the length of one 4 Hz theta cycle, there may be greater asymmetry in within-cycle connectivity on LSD and larger asymmetry in between-cycle connectivity on placebo.* *** $p < 0.001$, ** $p < 0.01$, * $p < 0.05$.

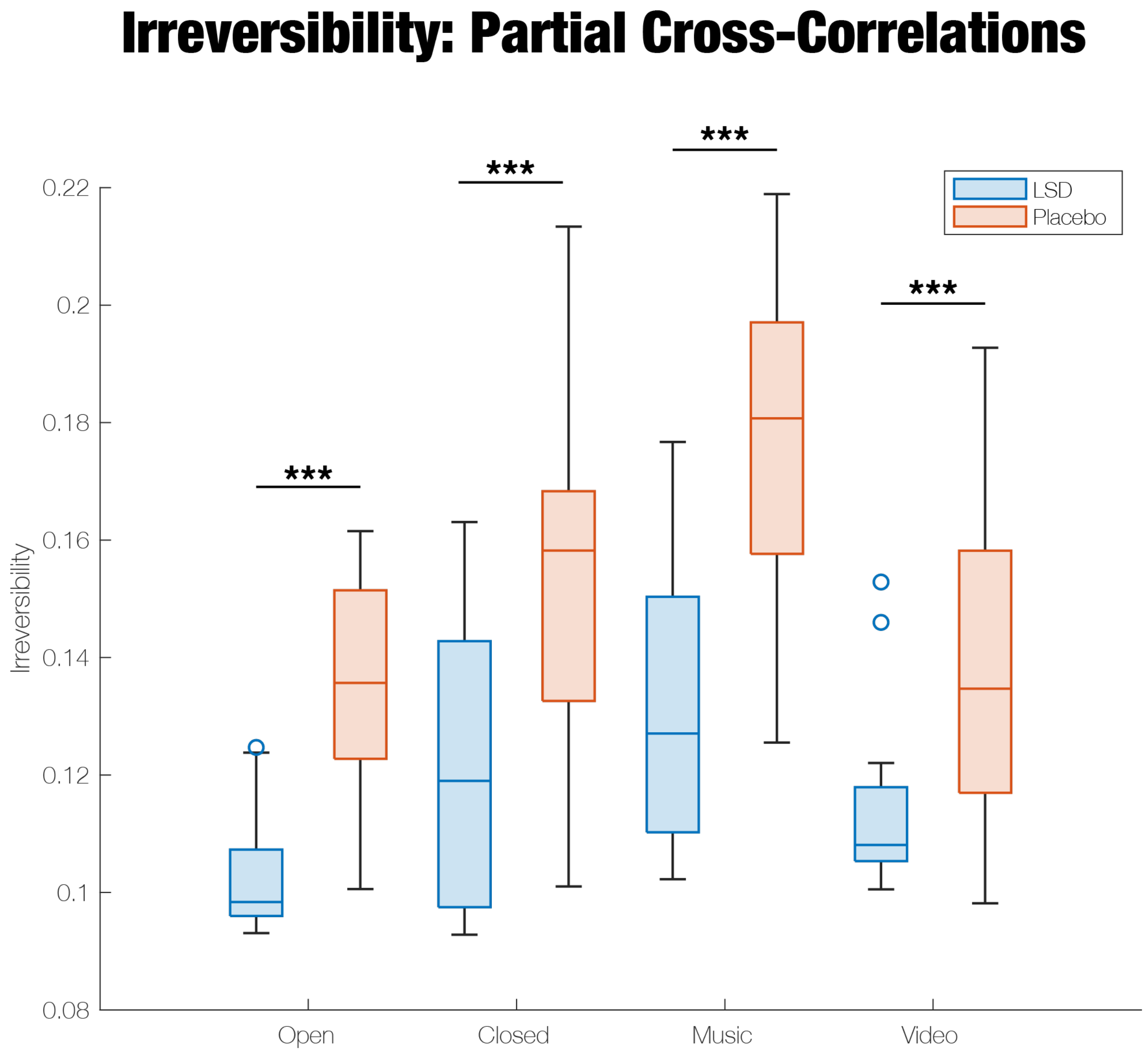


***Supplementary Figure 6. LSD significantly decreases irreversibility even with respect to partial cross-correlations rather than ordinary cross-correlations.*** *One limitation of the ordinary pairwise cross-correlations implemented in INSIDEOUT is that they do not control for confounding influences from other regions. This limitation can be resolved with partial cross-correlations. Here, timeseries were shifted by the same* $\tau$ *used in the main text (0.225 seconds). Then, each pair of timeseries was regressed against all other timeseries, and the correlation between the residuals was computed. Across all four conditions, irreversibility was still significantly lower on LSD when we used this method.* *** $p < 0.001$, ** $p < 0.01$, * $p < 0.05$.

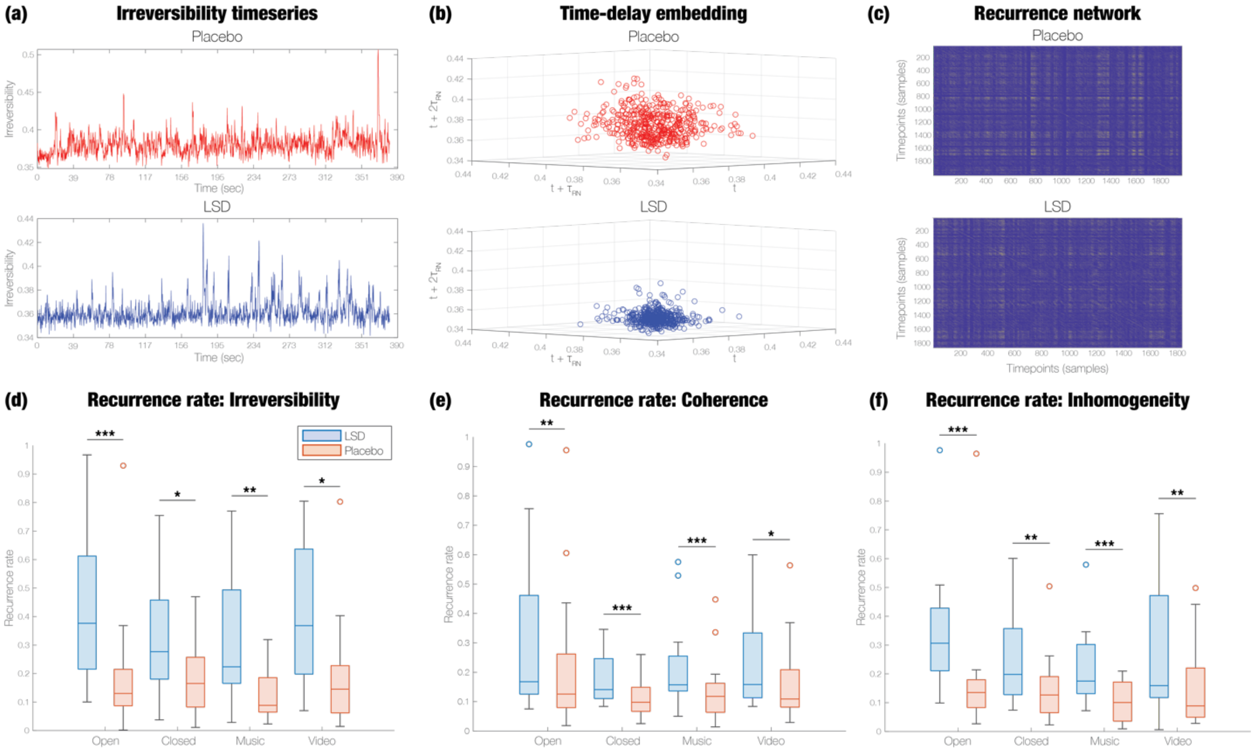


***Supplementary Figure 7. LSD constrains the dynamical repertoire of irreversibility, coherence, and inhomogeneity.*** *(a) We first performed a sliding-window analysis of each hierarchy metric in overlapping 1-second windows of data. Here, we display the irreversibility timeseries for one participant in the Open condition, under both placebo and LSD. (b) We conducted a time-delay embedding of each timeseries, where each dimension corresponds to the value of the metric across successive intervals of time. We show a 3-dimensional embedding of the same timeseries displayed in (a). LSD clearly reduces the density of points in the embedding. (c) We created a recurrence network, in which timepoints are connected by an edge if their distance in the embedding is below a certain threshold. Yellow entries correspond to the presence of an edge, and blue to the absence of an edge. Clearly, there are many more edges in the placebo recurrence network than the LSD recurrence network. (d-f) Across all participants and conditions, LSD significantly increases the recurrence rate of irreversibility, coherence, and inhomogeneity, indicating that the dynamics of each hierarchy metric are more similar over time under LSD compared to placebo.* *** $p < 0.001$, ** $p < 0.01$, * $p < 0.05$.

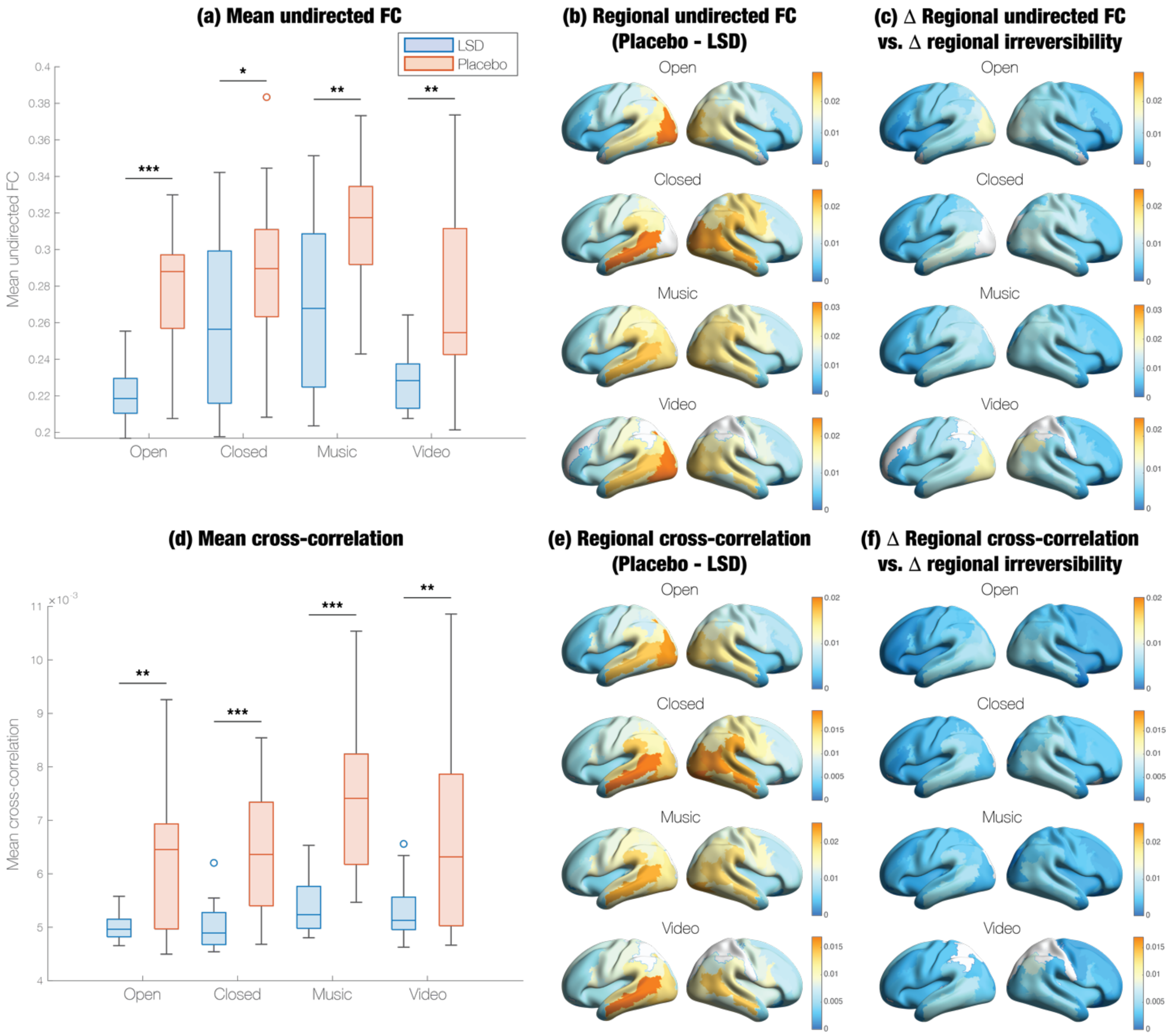


***Supplementary Figure 8. LSD's effect on regional irreversibility is highly similar to its effect on undirected functional connectivity and cross-correlations.*** *The left column (a, d) shows the mean across regions of each metric, the middle column (b, e) the regions that significantly differed in each metric between placebo and LSD, and the right column (c, f) the difference between regional changes in each metric and regional changes in irreversibility on LSD. Orange colours represent large changes and blue colours denote small changes. Each metric, both within nearly all regions and when averaged across regions, was significantly lower on LSD than placebo, in all four conditions. The spatial topographies of undirected FC (c) and time-forwards cross-correlations (f) are very similar to those of regional irreversibility.*

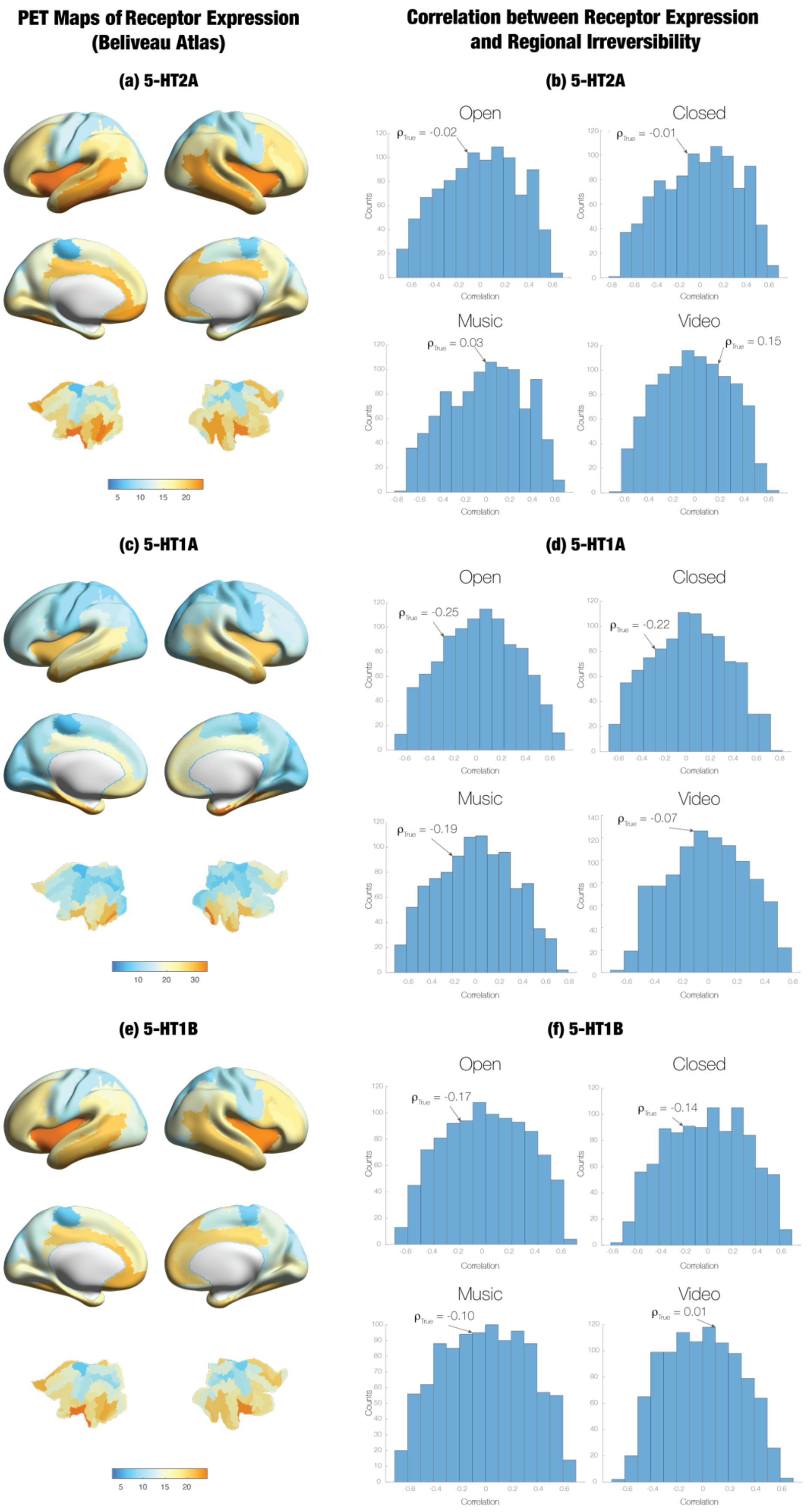
PET Maps of Receptor Expression
(Beliveau Atlas)
Correlation between Receptor Expression
and Regional Irreversibility
(a) 5-HT2A
(b) 5-HT2A
Open
ρTrue = -0.02
Closed
ρTrue = -0.01
Music
ρTrue = 0.03
Video
ρTrue = 0.15
(c) 5-HT1A
(d) 5-HT1A
Open
ρTrue = -0.25
Closed
ρTrue = -0.22
Music
ρTrue = -0.19
Video
ρTrue = -0.07
(e) 5-HT1B
(f) 5-HT1B
Open
ρTrue = -0.17
Closed
ρTrue = -0.14
Music
ρTrue = -0.10
Video
ρTrue = 0.01
Counts
Correlation

***Supplementary Figure 9. 5-HT$_{2A}$ concentration does not correlate with changes in regional irreversibility under under LSD.*** *LSD's agonism of the 5-HT$_{2A}$ receptor is crucial for its hallucinogenic and behavioural effects; if the receptor is blocked, then participants do not experience the "trip" (González-Maeso et al., 2007; Vollenweider et al., 1998). (a) Using PET data on 5-HT$_{2A}$ receptor expression in the human brain (Beliveau et al., 2017), we quantified the average concentration of the 5-HT$_{2A}$ receptor in each region of the AAL-90 parcellation, weighted by the probability that each voxel in each region was in grey matter. (Note: we also used other PET atlases, such as the Savli and Hansen atlases, and achieved similar results.) (b) We then correlated differences in regional irreversibility between LSD and placebo with the average regional concentration of 5-HT$_{2A}$ receptors, and we determined the significance of this correlation with respect to a permutation of the 5-HT$_{2A}$ distribution that preserved its spatial autocorrelation. There was no correlation in any of the four conditions. (c) and (e) show the regional density of the 5-HT$_{1A}$ and 5-HT$_{1B}$ receptor, respectively, based on the same PET maps. (d) and (f) display the correlation between 5-HT$_{1A}$ / 5-HT$_{1B}$ expression and regional irreversibility, with respect to the null distribution. Once again, there were no significant correlations.*

| *F*-test | | | | Post-hoc Tukey tests | | | | |
|---|---|---|---|---|---|---|---|---|
| Effect | DF | *F* | *p* | Condition | *p* | *T* | CI (upper) | CI (lower) |
| Drug | 1,15 | 46.925 | $5.50*10^{-6}$ | Open | $1.25*10^{-6}$ | -7.76 | -0.0067 | -0.0038 |
| Condition | 1.67,25.12 | 31.388 | $4.32*10^{-7}$ | Closed | $1.72*10^{-5}$ | -6.19 | -0.0087 | -0.0043 |
| Interaction | 3,45 | 6.585 | $8.74*10^{-4}$ | Music | $1.62*10^{-6}$ | -7.60 | -0.0110 | -0.0062 |
| | | | | Video | 0.0032 | -3.50 | -0.0077 | -0.0019 |

***Supplementary Table 1. Irreversibility statistics ($\tau$ = 0.225 seconds).*** *These statistics correspond to the results shown in Figure 2a/2b. Note that CI (upper) and CI (lower) correspond to the confidence interval of LSD – placebo.*

| *F*-test | | | | Post-hoc Tukey tests | | | | |
|---|---|---|---|---|---|---|---|---|
| Effect | DF | *F* | *p* | Condition | *p* | *T* | CI (upper) | CI (lower) |
| Drug | 1,15 | 33.011 | $3.87*10^{-5}$ | Open | $6.43*10^{-6}$ | -6.76 | -0.0021 | -0.0011 |
| Condition | 1.75,26.2 | 17.229 | $3.15*10^{-5}$ | Closed | 0.0002 | -4.99 | -0.0024 | -0.0010 |
| Interaction | 1.85,27.77 | 2.675 | 0.09 | Music | $1.02*10^{-5}$ | -6.49 | -0.0031 | -0.0016 |
| | | | | Video | 0.0062 | -3.18 | -0.0027 | -0.0005 |

***Supplementary Table 2. Hierarchical coherence statistics.*** *These statistics correspond to the results shown in Figure 3c/3d. Note that CI (upper) and CI (lower) correspond to the confidence interval of LSD – placebo.*

| *F*-test | | | | Post-hoc Tukey tests | | | | |
|---|---|---|---|---|---|---|---|---|
| Effect | DF | *F* | *p* | Condition | *p* | *T* | CI (upper) | CI (lower) |
| Drug | 1,15 | 22.712 | 0.0003 | Open | $7.43*10^{-6}$ | -6.67 | -0.0017 | -0.0009 |
| Condition | 1.83,27.39 | 13.311 | 0.0001 | Closed | 0.0139 | -2.78 | -0.0016 | -0.0002 |
| Interaction | 1.9,28.46 | 1.985 | 0.158 | Music | 0.0003 | -4.73 | -0.0020 | -0.0008 |
| | | | | Video | 0.0122 | -2.85 | -0.0015 | -0.0002 |

***Supplementary Table 3. Hierarchical inhomogeneity statistics.*** *These statistics correspond to the results shown in Figure 3e/3f. Note that CI (upper) and CI (lower) correspond to the confidence interval of LSD – placebo.*
